# Neuronal synchronization in *Drosophila*

**DOI:** 10.1101/2024.09.19.613913

**Authors:** Florencia Fernandez-Chiappe, Marcos Wappner, Luis G. Morelli, Nara I. Muraro

## Abstract

Rhythms are intrinsic to biological processes across temporal and spatial scales. In the brain, the synchronized oscillatory activity of neurons creates collective rhythms that are essential for complex functions. While this is a recognized phenomenon in the mammalian brain, information about insect neuronal synchrony and its underlying mechanisms is scarce. In the fly brain, neuronal oscillations were reported in individual lateral ventral neurons (LNvs), which play a key role in circadian and sleep behaviors. However, it is still unclear whether and how these participate in a collective rhythm. In this work, we perform thorough whole-cell patch clamp recordings of LNvs, and demonstrate consistent membrane potential oscillations. We show that oscillations degrade over time, and disappear upon exposure to an acetylcholine receptor blocker. Together with a flat phase response curve, these results suggest that oscillations are exogenously produced. Prompted by these results, we propose a generic forced oscillator theory that can account for the experimental phase response. The theory further predicts that neurons with similar properties should oscillate in synchrony with zero lags, while neurons with different properties may show coherent oscillations with non-zero lags. We confirm this prediction through simultaneous patch clamp recordings of neuronal pairs, revealing that large LNvs are consistently advanced relative to small LNvs. Additionally, we find that other neurons in the accessory medulla also exhibit coherent membrane potential oscillations, with diverse lags. Our findings suggest the intriguing possibility that brain waves may arise from collective neuronal activity within this region of the fly brain.

## INTRODUCTION

Collective rhythms, in the form of synchronized oscillations or propagating waves, pervade biological functions [1], such as molecular signaling in cell fate decisions [2], embryonic patterning [3], heart rhythms [4], animal motion control [5, 6], and population phenomena [7–9]. Synchronized neuronal oscillatory activity produces brain rhythms that underpin complex brain phenomena [10]. Rhythms of different spectral content have distinct roles in mammals. For instance, memory performance correlates with theta phase modulation of gamma rhythms, that cause cell assembly segregation in the hippocampus and entorhinal cortex of rats and humans [11]. Another example is brain rhythms measured by electroencephalography, that can be used to define sleep states [12, 13]. In the fly, neuronal oscillatory activity gives rise to brain rhythms as well, and these rhythms have been associated with different behavioral states. For example, selective attention to visual stimuli was linked to local field potential oscillations in the beta range, measured in the fly central complex [14]. Moreover, local field potential recordings revealed alpha oscillations in the central brain associated to a transitional sleep stage [15], and delta band rhythms in the ellipsoid body were associated with sleep behavior [16].

Neuronal oscillations are also present in lateral ventral neurons (LNvs) of *Drosophila* [17–19], a group of 16-18 circadian clock cells characterized by the expression of the neuropeptide Pigment Dispersing Factor (PDF) [20, 21]. LNvs somata are localized in the accessory medulla (aMe) and extend their axonal projections beyond this region [22, 23]. LNvs are further subdivided in two groups according to their size, anatomy, function and developmental history. The 8 small-LNvs (sLNvs) possess small-sized somas and together constitute a hierarchical circadian pacemaker under free running conditions [24–27]. The 8-10 large-LNvs (lLNvs) have more sizable somas and have been characterized as arousal neurons [28–31]. A previous study reported that sLNvs and lLNvs membrane potential oscillate with a similar frequency [17]. However, whether these neuronal oscillations are part of a collective rhythm is still unknown.

Here, we perform whole-cell patch clamp electrophysiological recordings in *Drosophila* neurons, and develop a quantitative analysis pipeline to characterize membrane potential oscillations. We show that LNvs oscillations become slower over time and depend on cholinergic neurotransmission. Together with a flat phase response, this suggests that oscillations are not cell-autonomous. We then propose a generic forced oscillator theory which captures the experimental observations. The theory further predicts that synchronized oscillations should occur with zero lags between neuronal pairs of the same type, while non-zero lags may occur between oscillations of different neuronal types. We confirm this prediction by means of simultaneous patch clamp experiments, revealing that lLNv oscillation is consistently advanced with respect to the sLNv. Finally, we discover that other neurons in the accessory medulla show membrane potential oscillations as well, and that these are coherent with LNv oscillations. Altogether, our findings open up the exciting possibility of brain waves resulting from collective neuronal activity present in this relatively unexplored region of the fly brain.

## RESULTS

### Consistent oscillations in individual LNvs

*Drosophila* LNvs fire action potentials that are organized in bursts, a firing mode that has previously been associated with neuropeptide release [32–35]. In this firing modality, action potentials are mounted on membrane depolarization oscillations. To characterize LNv oscillations, we performed extensive whole-cell patch clamp recordings of LNvs in brain explants following previously established methods [36–38] (Fig. 1A). Recordings were done during the light phase of flies entrained to 12:12 light:dark cycles. Experiments were done in Pdf-RFP female flies [39] where the pigment dispersing factor gene promoter directly drives the expression of the Red Fluorescent Protein. LNvs were identified by the genetically encoded fluorescent marker, their anatomical position and size. Selected neurons were accessed into whole-cell patch clamp configuration in voltage-clamp mode, and promptly switched to current-clamp (with I=0) for the recording of membrane voltage. In this way, we obtained long electrophysiological recordings displaying clear cycles of membrane potential (Fig. 1B). To quantitate these oscillations, we first applied filters that remove fast and slow frequencies from the raw data (Methods, Fig. S1), smoothing out the high frequency noise and spiking activity, and removing slow trends present in some of the long recordings, without distorting the cycles shape (Fig. 1C). Next, we defined cycle duration (CD) as the time between two equivalent points in the voltage oscillation cycle. We found that CD was most reliably determined by taking a threshold crossing of the cycle upstroke, which has the steeper slope, and averaging over a range of threshold values (Fig. 1C). We examined the stability of CD across recordings ranging from 2 to 46 minutes in duration (Fig. 1D), comprising between 18 to 748 cycles. While lLNvs oscillations appear quite regular, a cycle is occasionally skipped when the signal does not reach the threshold, resulting in a spurious cycle that doubles the average CD (red dots in Fig. 1C, E and Fig. S2). We excluded those anomalies from the analysis, rejecting the CD values that were outside the general trend in a two step process, the remaining 97% of cycles were labeled as valid (Methods).

**Figure 1.**
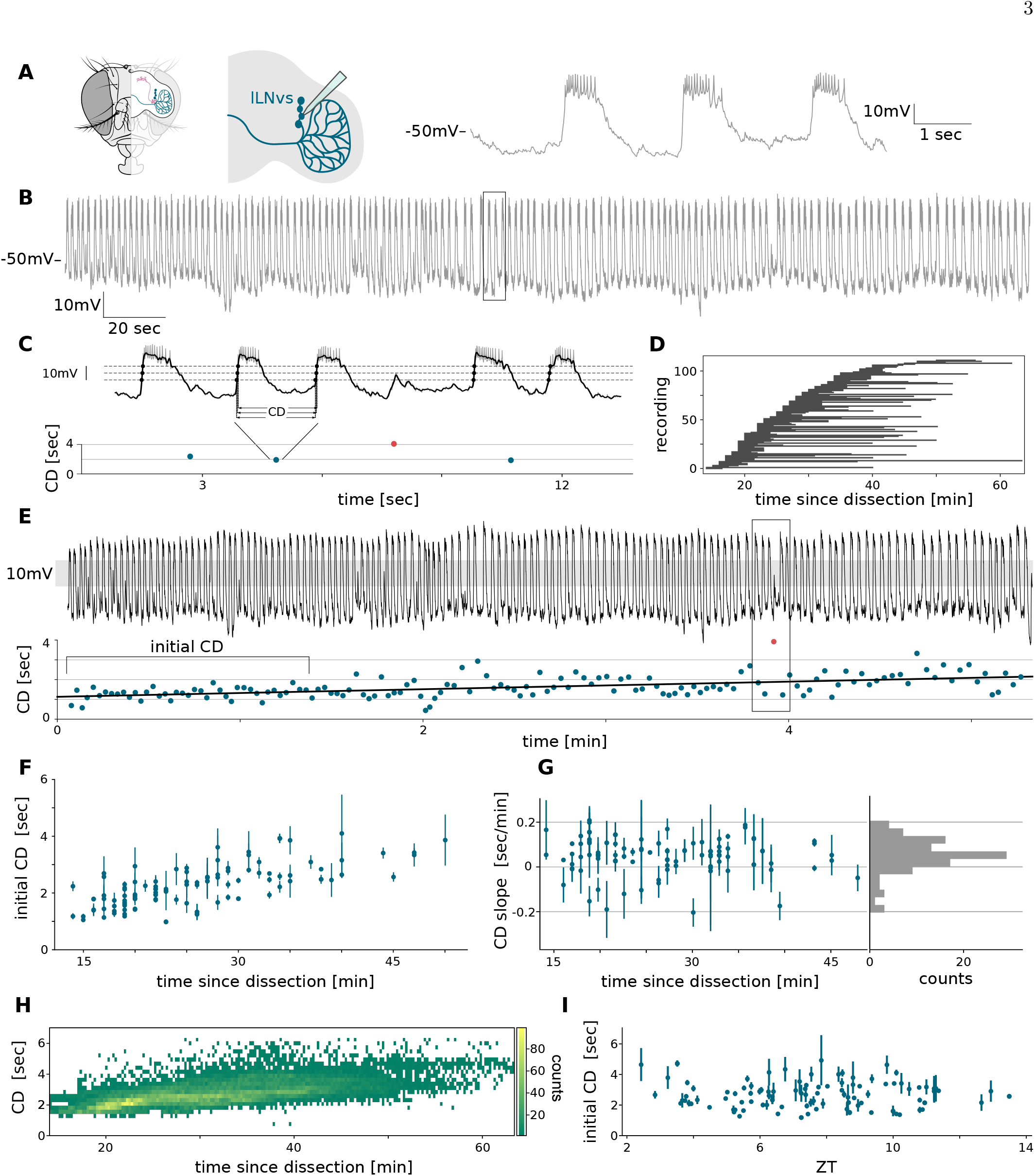
lLNvs display membrane potential oscillations. (A) Left: Scheme of the *Drosophila* head. Middle: Scheme of brain explant (only right side is shown) displaying lLNvs and recording electrode. Right: zoom into the box from B. (B) Representative recording from a lLNv, beginning 22 min after dissection. (C) Top: fraction of detrended recording from B (grey line) and denoised trace (black line). Three threshold values (dashed lines) and corresponding threshold crossings (black dots) are indicated. Bottom: corresponding valid (blue dots) and non valid (red dot) CD measurements. (D) Duration of *n* = 112 individual recordings represented by horizontal bars. (E) Top: detrended and denoised recording from B. Shaded region indicates the threshold range. Bottom: corresponding CD measurements (color dots) and linear fit (line). Box marks the zoom in C and bracket indicates the span for initial CD calculation. (F) Initial CD vs. time since dissection (dots) are positively correlated (Pearson *r* = 0.67). Error bars are the standard deviation (SD). (G) Left: CD slope vs. time since dissection (dots) show no correlation (Pearson *r* = 0.22). Error bars are slope fitting errors. Right: histogram of CD slopes, mean is 0.09 sec/min. Slope is significantly different from zero (*p <* 10^*−*8^, Wilcoxon signed-rank test). (H) 2D histogram of pooled CD values (24949 cycles) from all lLNvs recorded, aligned in time since dissection. (I) Initial CD vs. ZT show no correlation (Pearson *r <* 10^*−*1^). Error bars are SD.

Previous reports showed that LNvs membrane oscillation frequency decays with time in ex vivo preparations [17, 18]. Therefore, we registered the time since dissection, from the moment of brain removal, through the ex vivo explant preparation [36] to the start of voltage data acquisition for each recording [37]. In agreement with previous reports, here we observe that in individual recordings there is a systematic increase of CD as a function of time since dissection (Fig. 1E). This trend is also reflected globally at the population level, where the initial CD, defined as an average of the first 40 valid cycles, increases with the time since dissection for most recordings (Fig. 1F). Consistently, the slopes of the trendlines obtained from fitting the CD for individual recordings (illustrated in Fig. 1E, bottom panel, black line) are predominantly positive and independent of time since dissection (Fig. 1G). The increase of the CD as a function of time since dissection is clearly visible in a heatmap representing the CD values of all recorded cells (Fig. 1H). Contrary to the CD, we found that the baseline potential (the trough value of the oscillation) and the amplitude (peak to trough) of the recordings do not follow a general trend as a function of time since dissection (Figs. S3 and S4). lLNvs are clock neurons, and their activity has been reported to change between day and night [18, 40, 41]. For this reason and with the aim of reducing variability, we restricted our experiments to the light phase of the diel cycle. During this phase, the initial CD lacked a clear trend as a function of the time of day (Zeitgeber Time, ZT) (Fig. 1I), therefore ZT was disregarded as a variable for further analysis.

Electrophysiological recordings of sLNvs have been scarcely reported in the literature [17, 40, 42] probably due to their small size, which represents a challenge for the technique. In spite of this, we have accomplished a thorough physiological analysis of these hierarchical clock neurons as oscillators. Similar to lLNvs, sLNvs present oscillations in their membrane potential (Fig. 2A). We examined the stability of CD in sLNv recordings with durations spanning from 2 to 17 minutes (Fig. 2B), and found a systematic increase of CD as a function of time since dissection in this neuronal group as well. Similar to the lLNvs, this phenomenon can be appreciated in individual recordings (Fig. 2C) and at the population level (Fig. 2D, E, F). Also for sLNvs, the initial CD lacked a clear trend as a function of ZT during daylight (Fig. 2G). In summary, our quantitative characterization of LNvs membrane potential shows that both, lLNvs and sLNvs, display consistent oscillations with a cycle duration that increases gradually with time ex vivo.

**Figure 2.**
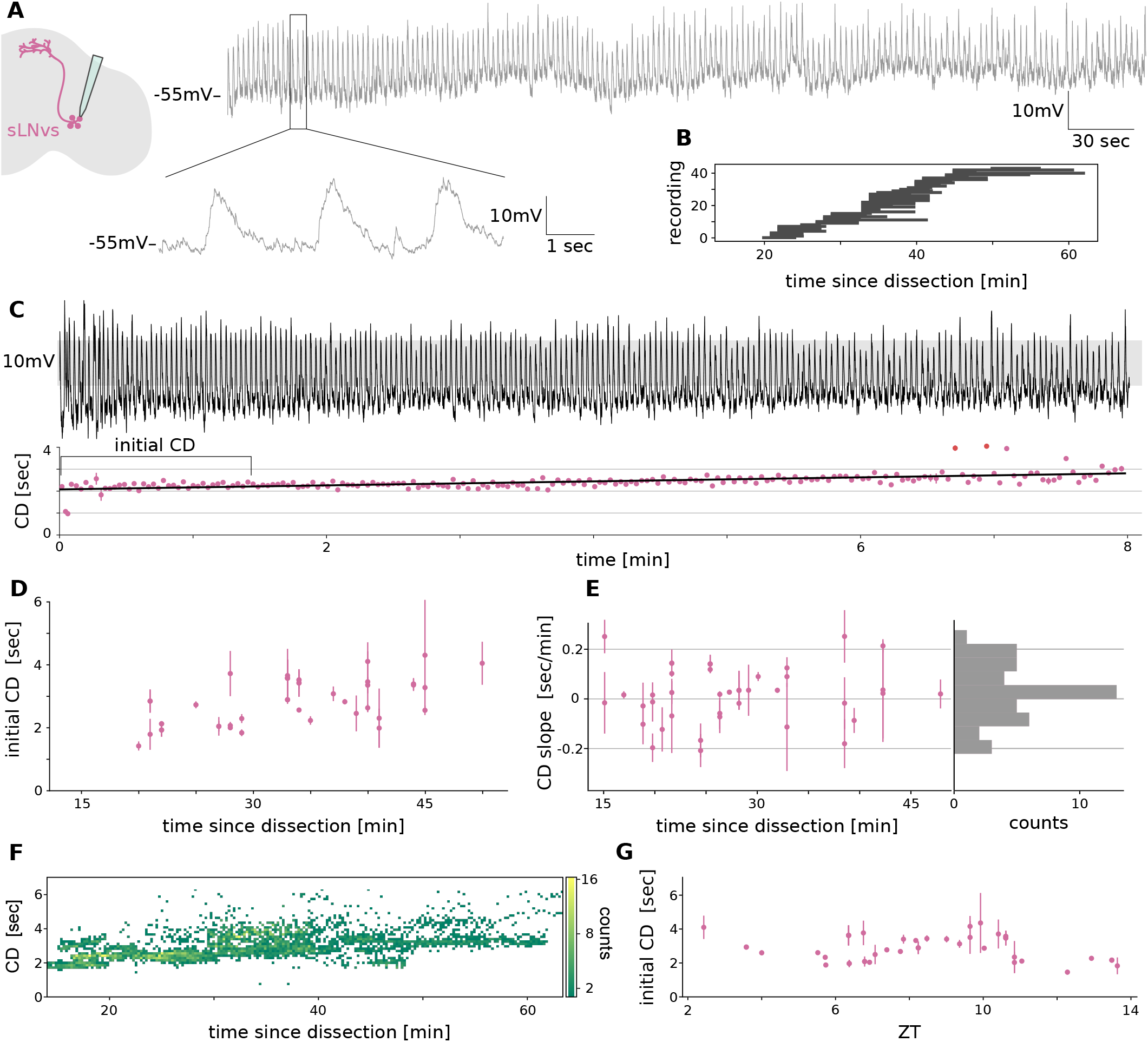
sLNvs display membrane potential oscillations. (A) Left: scheme of brain explant displaying sLNvs and recording electrode. Right: representative recording from a sLNv, beginning 35 min after dissection. Inset: zoom into the box. (B) Duration of *n* = 44 individual recordings represented by horizontal bars. (C) Top: detrended and denoised recording from A. Shaded region indicates the threshold range. Bottom: corresponding valid (pink dots) and non valid (red dots) CD measurements, and linear fit (line). Bracket indicates the span for initial CD calculation. (D) Initial CD vs. time since dissection (dots) are positively correlated (Pearson *r* = 0.62). Error bars are SD. (E) Left: CD slope vs. time since dissection (dots) show no correlation (Pearson *r* =*−*5 *×* 10^*−*3^). Error bars are slope fitting errors. Right: histogram of CD slopes, mean is 0.021 sec/min. Slope is significantly different from zero (*p <* 0.5 *×* 10^*−*1^, Wilcoxon signed-rank test). (F) 2D histogram of pooled CD values (5001 cycles) from all sLNvs recorded, aligned in time since dissection. (G) Initial CD vs. ZT show no correlation (Pearson *r <* 10^*−*1^). Error bars are SD.

### LNv oscillations are exogenously driven

What can we learn about LNvs biology from the CD increase observed in ex vivo preparations? In theory, oscillations in neuronal membrane potential may be cell-autonomous, driven by the sequential action of specific ion channels [43], or they could be driven by synaptic inputs. An example of a cell-autonomous mechanism for membrane potential oscillation is the one relying on a hyperpolarization-activated current such as I_h_ and the T-type low-voltage activated calcium channel [44], that together orchestrate self-sustained membrane potential oscillations. We would expect that, if lLNvs oscillatory activity is cell-autonomous, the phase of the oscillation would be shifted by the injection of current directly into the cell, which is expected to affect the gating mechanisms of voltage-dependent channels [43].

To test this hypothesis, we constructed a phase-response curve (PRC) [45]. We evaluated how lLNv oscillations were affected by the injection of short pulses of negative current, delivered randomly at different times relative to the start of the oscillation cycle (Fig. 3A).

**Figure 3.**
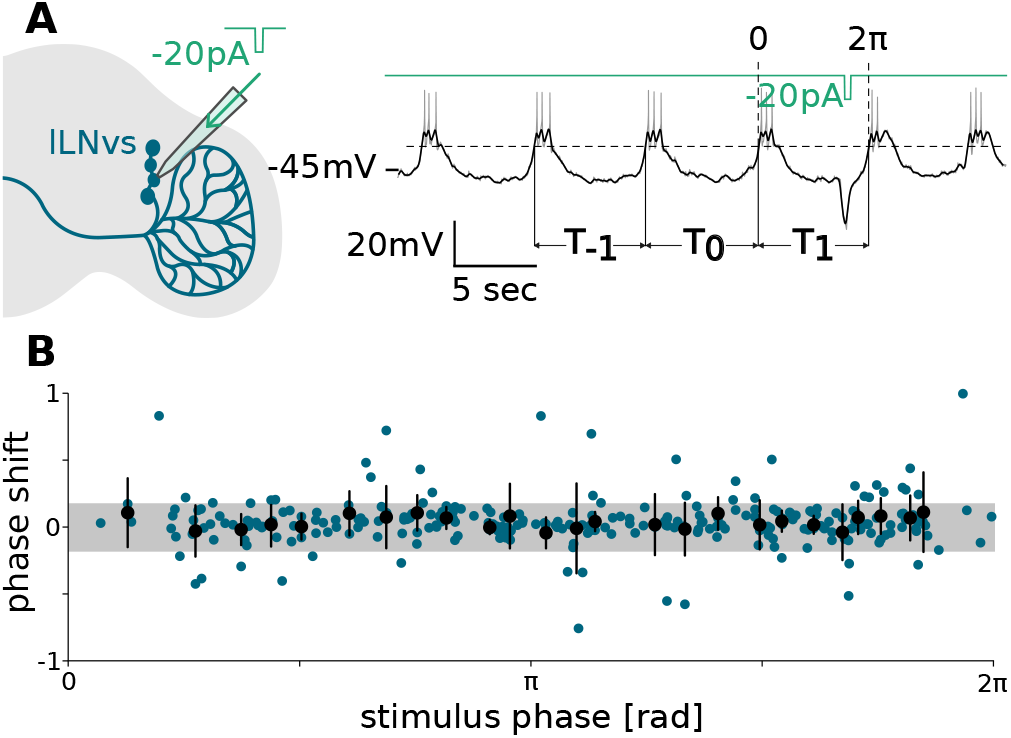
Phase response curve is flat. (A) Left: scheme of brain explant displaying lLNvs, electrode and current injection. Right: recording fragment from current injection experiment (grey line), denoised trace (black line), and injected current (green line). Representative threshold (horizontal dashed line) used to define the perturbed and two previous cycles as indicated, and the mapping of *T*_1_ onto the [0, 2*π*] interval are shown. (B) Phase shift *P*_1_ as a function of stimulus phase (Methods). Blue dots correspond to a single perturbed cycle, from *n* = 10 different neurons, totaling 274 cycles. Black dots are average of 11 consecutive blue dots, and error bars are SD. Shaded area is SD of the reference cycles.

We defined a phase shift as the difference between the CD of the perturbed cycle and the previous unperturbed cycle, relative to the latter (Methods). The resulting phase shifts caused by perturbations appeared noisy and did not follow a clear function of the stimulus phase (Fig. 3B). As a control, we compared these phase shift values with the naturally occurring variability. We defined this variability as the standard deviation of the relative difference between the CD of the two previous unperturbed cycles (shaded area in Fig. 3B, Methods). We observed that about 80% of measurements fall within this variability range. Thus, we conclude that the PRC is flat, suggesting that there is no strong autonomous component to the LNv oscillations.

An alternative hypothesis is that LNvs rely on exogenous information to produce these oscillations of membrane potential. lLNvs projections are located throughout the optic lobes, where acetylcholine (ACh) is the main neurotransmitter [46] prompting us to test its involvement with lLNv membrane oscillations. lLNv membrane oscillations were abolished by treatment with the nicotinic ACh receptor antagonist mecamylamine (mec) (Fig. 4A). Within minutes after mec exposure on the preparation bath, lLNv oscillations were invariably lost (*n* = 12), action potentials stopped their organization in bursts and firing became tonic and sparser, consistent with previous observations [18, 47]. We next tested whether sLNvs oscillations were similarly dependent on ACh, which has not been reported before. Upon exposure to mec, we found that sLNvs also lost their oscillatory activity (Fig. 4B, *n* = 14).

**Figure 4.**
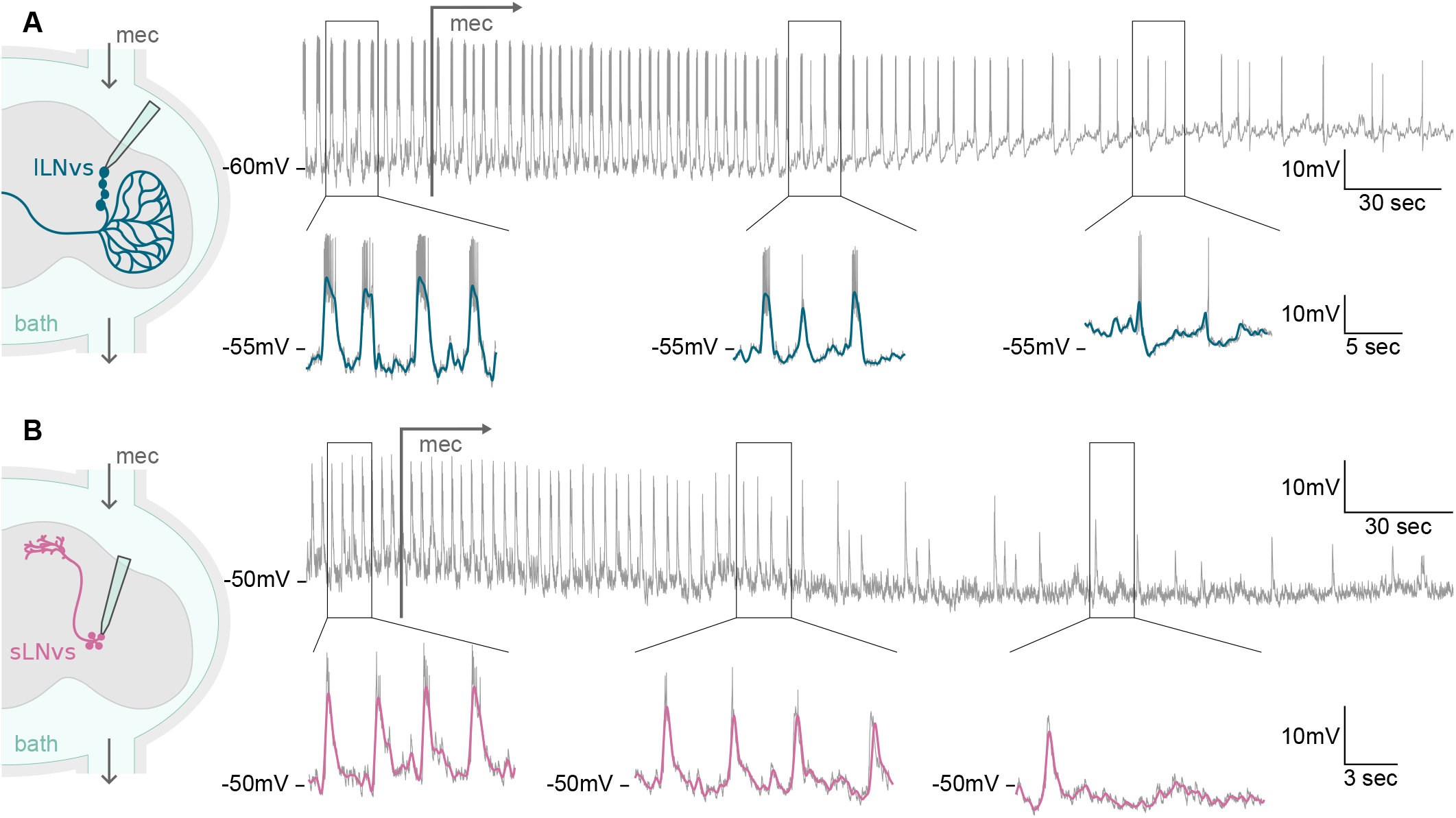
Membrane oscillations are ACh-dependent. (A, B) Left: scheme of brain explant in a bath with mec perfusion, displaying (A) lLNvs and (B) sLNvs, and recording electrode. Right: Representative recordings of mec treated neurons, with gray arrow indicating the beginning of mec application. Insets show zoom into boxes displaying fragments of the recording (grey line) and denoised traces of (A) lLNvs (blue line) and (B) sLNvs (pink line). Recordings performed (A) 56 min and (B) 37 min after dissection.

In summary, we observed that LNvs membrane oscillations were altered in two contrasting ways. On the one hand, neuronal oscillations are gradually slowed by the ex vivo preparation of the brain (Figs. 1 and 2). On the other hand, neuronal oscillations are sharply obliterated by the treatment with cholinergic blockers (Fig. 4). Together with the flat PRC (Fig. 3), these results indicate that LNvs membrane oscillations are not cell-autonomous, they depend on cholinergic circuits that are subtly disturbed during the brain preparation and result in the gradual decay of neuronal oscillations.

### Theoretical description of a forced oscillatory membrane potential

Next, we seek to gain insight into the conditions that could lead to a flat PRC (Fig. 3B). We introduce a generic theory for a forced oscillator that describes key aspects of the oscillatory dynamics without knowledge of the molecular and physiological underpinnings [48] (Methods). Furthermore, with different parameter values, the same theory can be used to describe diverse neuronal types. Dynamical systems theory reveals how oscillations can arise in a system when parameters change [49]. Here we focus on a mechanism common to many neuronal models [43], where a stable stationary state may lose stability giving rise to sustained autonomous oscillations, termed a Hopf bifurcation [49].

We describe the state of the system in polar coordinates using the phase *θ* and the amplitude *r* (Supplemental Methods). We introduce a harmonic forcing with strength *f* and frequency Ω, so the forcing phase *ψ* is

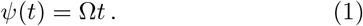

The dynamics of the forced oscillator is given by

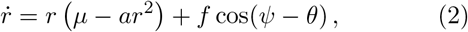

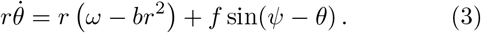

The value of *µ* determines whether the system presents sustained oscillations (*µ >* 0) or a single stationary state *r* = 0 (*µ <* 0) [49]. Parameter *a* influences how quickly the system returns to its steady state after a perturbation, *ω* is a characteristic frequency of the system, and *b* is an amplitude-dependent correction to this frequency. To formalize the hypothesis that the oscillations are externally driven we set *µ <* 0, such that the system does not sustain autonomous oscillations. When a forcing *f >* 0 is introduced, the oscillations turn on (Fig. 5A).

**Figure 5.**
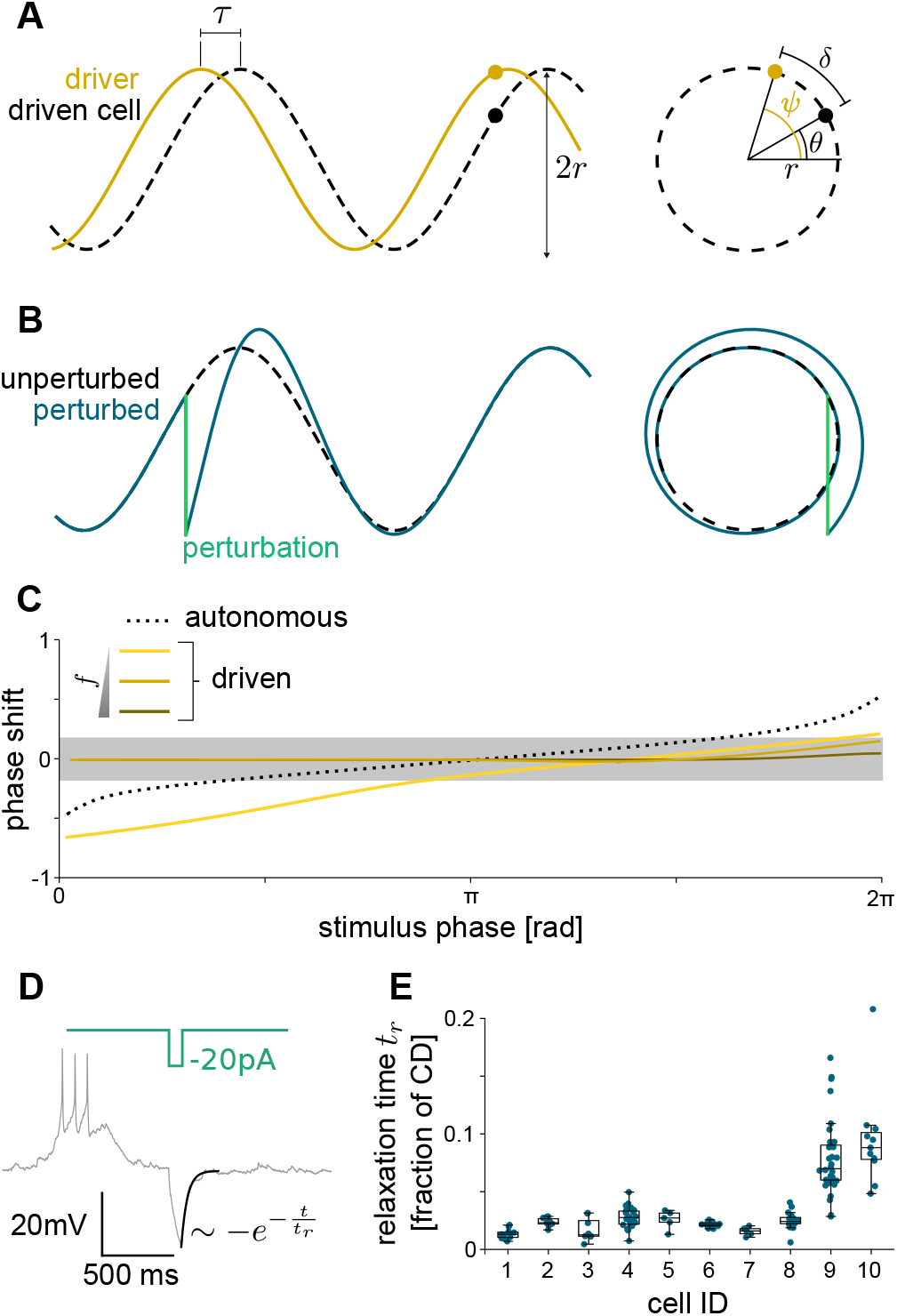
A forced oscillator theory is consistent with a flat PRC. (A) Left: numerical solutions to Eqs. (1) to (3) showing the driver (yellow line and dot) and the driven cell (dashed line and black dot). Time lag *τ* and amplitude 2*r* are indicated. Right: polar coordinates representation of the oscillations (dashed line), showing polar amplitude *r*, driver phase *ψ* (yellow dot), driven cell phase *θ* (black dot), and phase lag *δ*. Driver amplitude is not drawn to scale and parameters are chosen for illustration. (B) Numerical solution of an unperturbed oscillator (dashed line) and a perturbed oscillator (blue line). Instantaneous perturbation is indicated (green line). Left: temporal representation. Right: polar coordinates representation. (C) Numerical PRC of the model. Parameters as in Table I, except *f* = 0 and *µ* = 1 for autonomous oscillations (dashed line), and *f* = 104, 10.4, 1.04 for driven oscillations (darker to lighter yellow lines). (D) Fragment of recording from a current injection experiment (grey line), with exponential fit of the perturbation response (black line), and current injection (green line). (E) Relaxation time *t*_*r*_ scaled by CD at the time of perturbation *T*_1_ (dots) (Methods). Boxes are the interquartile range, bar is the median and whiskers extend 1.5 times the interquartile range.

To construct a theoretical PRC, we introduce perturbations in the model that mimic those of the experiment. We interpret the variable *y* = *r* sin *θ* as the membrane potential, and apply instantaneous shifts *δy* along the negative direction in *y* axis, at different points in the cycle (Fig. 5B). As expected, immediately after the perturbation, the phase is altered with respect to its unperturbed value. Then the forcing gradually returns the oscillation to steady state cycling. We compute a theoretical PRC in the same way we did for the experimental data: we measure the shift in cycle duration, normalized to the duration of the previous cycle.

**Table I.** Parameter and observable values used in the model.

| parameter | sLNv | sLNv | description |
| --- | --- | --- | --- |
| $\mu$ | -3.46 | -3.88 | bifurcation parameter |
| $a$ | $5.53/r_0^2$ | $5.37/r_0^2$ | limit cycle attractor strength |
| $\Delta$ | $-0.6 \Omega$ | $0.6 \Omega$ | detuning |
| $b$ | 0 | 0 | non-isochronicity |
| $f$ | $9.14 r_0$ | $9.43 r_0$ | forcing strength |
| observable | sLNv | sLNv | description |
| $\delta$ | $-0.03 \ 2\pi$ | $0.03 \ 2\pi$ | phase lag with forcing |
| $\rho_0$ | 11.4 | 8.85 | half-amplitude |
| $t_r$ | 0.023 | 0.023 | relaxation time |

For positive values of *µ*, we observe the typical shape of the PRC for autonomous oscillators near a Hopf bifurcation (Fig. 5C) [43]. For negative *µ* and positive forcing strength *f*, we find that the PRC becomes flatter for increasing *f* (Fig. 5C), until it falls within the uncertainty interval defined previously for the experiment. These large values of *f* would be accompanied by a fast return to the steady state, consistent with visual inspection of experimental PRC recordings (Fig. 3A). To quantitate this, we determined experimentally the characteristic decay time after the perturbation. We focused on the pulses that fall in troughs of the cycle, as these can be simply described by an exponential, and fitted an exponential relaxation after the current pulse was delivered (Fig. 5D, Methods). Characteristic relaxation times were consistently small with respect to cycle duration, *t*_*r*_ = (0.023 ± 0.008) CD, with two neurons showing slightly larger values, still below 0.1 CD (Fig. 5E). In summary, the theory can account for the flat phase response that we obtained experimentally, and characteristic relaxation times appear consistent with theoretical requirements.

### The theory predicts a phase lag with the forcing

Equations (1) to (3) describe the dynamics of amplitude *r*(*t*) and phase *θ*(*t*) of the forced oscillator. We next consider a steady state after transient perturbations elapse, in which the oscillator follows the driver with a constant amplitude *r*(*t*) = *r*_0_, that does not depend on time. In this steady state, the phase changes at the same frequency as the driving, so the instantaneous frequency is 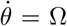, and *θ*(*t*) = Ω*t* + *δ*, where *δ* is a phase lag between the oscillator and the driver (Fig. 5A). This phase lag between the forced oscillator voltage and the forcing implies a time lag

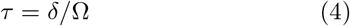

between the two signals. Replacing the steady state condition in Eqs. (1) to (3) we obtain expressions for the steady state amplitude and phase lag (Supplemental Methods):

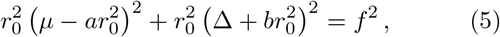

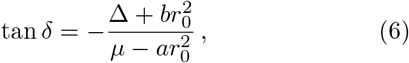

where the detuning Δ = Ω − *ω* is the difference between the driving frequency Ω and the characteristic frequency of the system *ω*. The lag *τ* between a neuron and the forcing is determined by parameters through Eqs. (4) to (6). We expect these parameters to have similar values among neurons of the same type, resulting in similar lags with respect to the forcing. Thus, the lag between neurons of the same type should be around zero. In contrast, different neuronal types may differ in their biophysical properties, resulting in distinct values for *τ*. Therefore, we expect neurons of different types to display a non-zero lag between them.

### LNvs display synchronized oscillations with a consistent lag

To test the predictions from the theory, we performed simultaneous patch clamp recordings of LNv pairs. These dual recordings are technically challenging because of the low success rate of the technique. As expected from the theory, LNv oscillations are coherent, that is, membrane potential cycles in the two cells are always coordinated throughout the recording (Fig. 6A). We observed that this occurs in all dual recordings for lLNv-lLNv pairs (Fig. 6B, *n* = 9), sLNv-sLNv pairs (Fig. 6C, *n* = 9) and sLNv-lLNv pairs (Fig. 6D, *n* = 18). This coherence is blatantly exposed when their usually very regular oscillatory pattern is briefly, and concomitantly, disrupted (Fig. S5A-C). Furthermore, upon exposure to mec, dual recordings show a correlated decay of the oscillatory dynamics (Fig. S5D, E).

**Figure 6.**
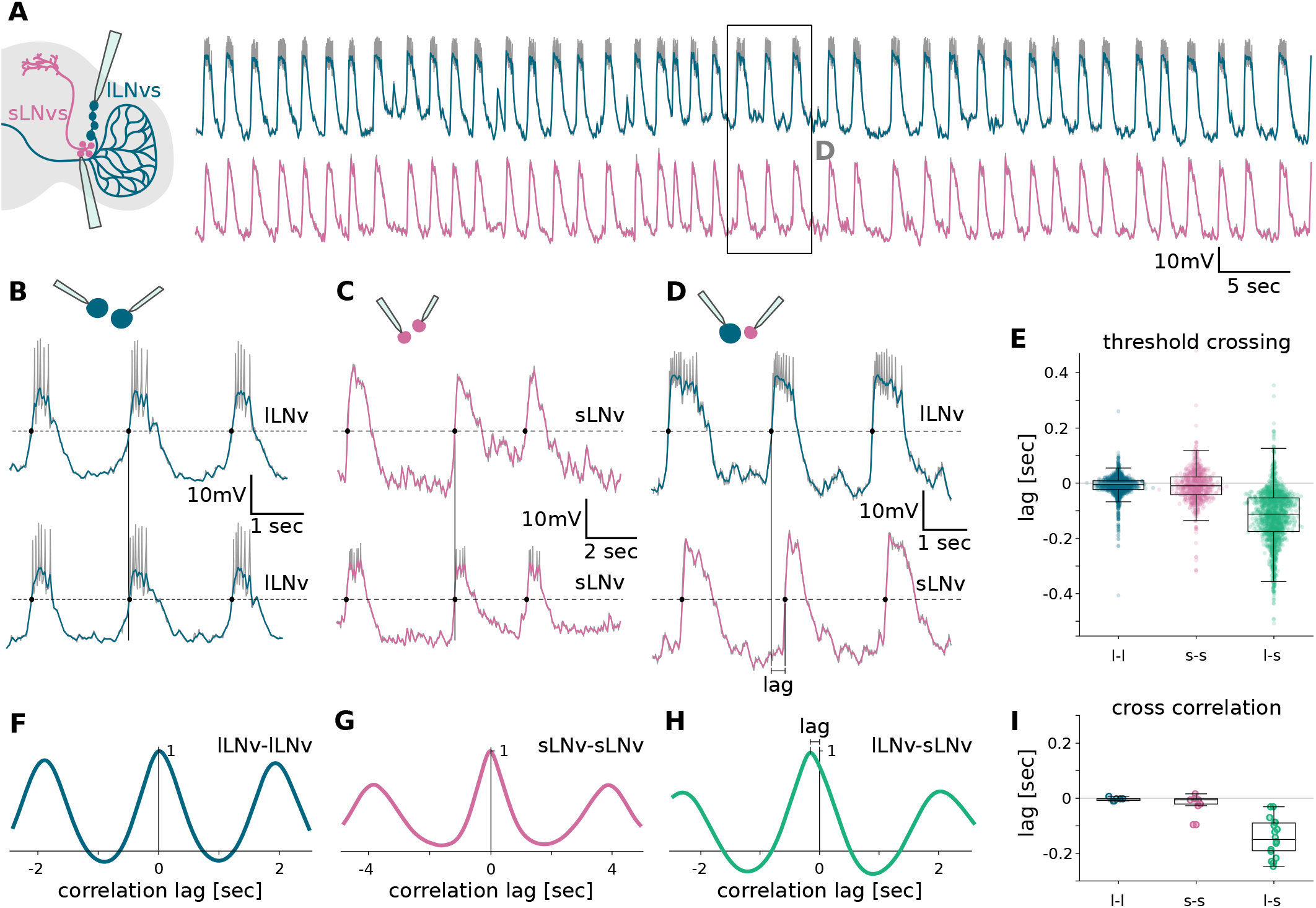
There is a lag between lLNvs and sLNvs membrane oscillations. (A) Left: scheme of brain explant displaying lLNvs and sLNvs together with recording electrodes. Right: Fragment of representative dual recording from a lLNv (blue) and a sLNv (pink), beginning 33 minutes after dissection. Box marks zoom in D. (B-D) Top: scheme of recorded (B) lLNv pairs, (C) sLNv pairs, and (D) lLNv-sLNv pairs. Bottom: Representative fragments of corresponding dual recordings with threshold crossing analysis (dashed line and dots). Vertical lines guide the eye to highlight the absence (B, C) or presence (D) of a time lag. In (A-D) data is detrended (grey) and further denoised (color). (E)Quantification of lag measurements from threshold crossing analysis for different LNv pairs, *n* = 9 lLNv-lLNv, *n* = 9 sLNv-sLNv and *n* = 18 lLNv-sLNv, totaling 819, 1708 and 1883 individual cycles (dots), respectively. (F-H) Representative cross correlation functions for (F) lLNv pairs (G) sLNv pairs and (H) lLNv-sLNv pairs. (I) Quantification of lags from cross correlation analysis. (E, I) Boxes are the interquartile range, bar is the median and whiskers extend to 1.5 times the interquartile range.

To quantify the degree of correlation between membrane oscillatory activity of neuronal pairs we computed differences between threshold crossing times, as previously introduced. Threshold crossing times were obtained for individual channels and subsequently matched between channels to find the closest crossing times, from which a time lag was obtained (Fig. 6B-E, Methods). This quantification revealed that in pairs of the same type there is no lag between threshold crossing times, while in pairs of different type, sLNvs consistently lag (0.12 ± 0.02) sec behind of lLNvs (Fig. 6E). To validate these results we introduced a cross-correlation analysis, as an orthogonal approach to determine the lag of the signals from our simultaneous patch clamp recordings (Methods). Using this methodology we obtained similar results, a high synchronization level of the oscillations of the same-cell pairs (Fig. 6F, G, I) and a lag of (0.14 ± 0.03) sec with the lLNv advanced with respect to the sLNv for the different-cell pairs (Fig. 6H, I). The same trends could be visualized on the individual dual recordings for both methods (Fig. S6 A, B), with a relatively constant lag between cells as time ex vivo progresses (Fig. S6 C, D).

In summary, our results show high synchronization levels of membrane oscillations in pairs of LNvs. While neurons of the same type oscillate in synchrony without a lag, neurons of different types show a significant lag. This is consistent with the theory, where the lags are interpreted as a consequence of differences in the parameters that define neuronal types.

### The theory can account for experimental observations of LNv membrane oscillations

The steady state condition provides relations between parameter values and observables such as the amplitude and phase lag of the forced oscillations, Eqs. (5) and (6). Additionally, we can obtain an expression for the relaxation time *t*_*r*_ from linear stability analysis [49]. Given that we have measured some of these observables, these relations set constraints that have to be satisfied by parameter values. This means that by choosing the values for some of the parameters and missing observables, the rest becomes uniquely determined.

Here we choose to vary the detuning Δ and phase lag *δ*, and determine the values of *µ, a* and *f* for small and large LNvs. The reason for this choice is that, although we do not know the precise values of these quantities, we can make an informed guess for their ranges. On one hand, we reasoned that the characteristic frequency of lLNvs and sLNvs might be similar to that of the neurons that drive the oscillations, that is *ω/*Ω∼ 1. On the other hand, we know the relative phase lag between lLNvs and sLNvs from experiments (Fig. 6). We use the value of this relative phase lag as a reference scale for phase lag values with respect to the forcing.

We scale the parameters so that they have the same units and we can compare their corresponding contributions in Eqs. (2) and (3). We find that it is possible to choose a set of values for both sLNvs and lLNvs that is consistent with all constraints arising from experimental data (Fig. 7A). To accommodate for the relative lag between LNvs, we choose parameters such that lLNvs are faster and advanced with respect to the forcing, and sLNvs are slower and delayed (Fig. 7A). With this parameter choice, scaled parameter values are similar, indicating that all terms contribute similarly to the dynamics, with a slightly stronger forcing term. Additionally, the values of *µ, a* and *f* are similar for both LNv types.

**Figure 7.**
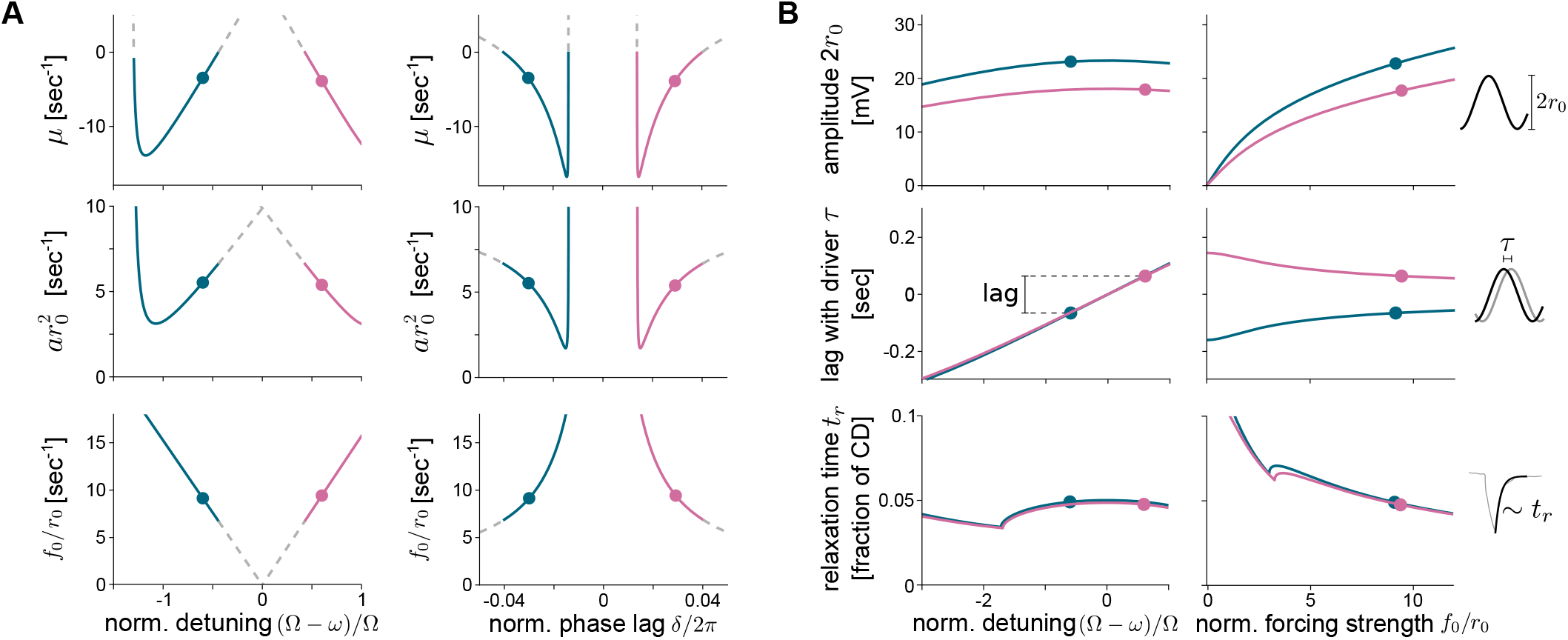
Theoretical relations between parameters and observables. (A) Parameter relations Eqs. (45-47) for lLNvs (blue line) and sLNvs (pink line) with parameters and observables as in Tab. I (dots). Gray dashed lines indicate parameter choices that result in *µ >* 0. Parameters *a* and *f* are scaled using *r*_0_ to have the same units as *µ*, so that their magnitudes can be compared. Phase lag *δ* is normalized to 2*π*, and detuning Δ = Ω *− ω* to the forcing frequency Ω. (B) Amplitude 2*r*_0_, time lag *τ* and relaxation time *t*_*r*_ scaled by CD, as a function of normalized detuning and forcing strength *f* normalized by amplitude *r*_0_. Vertical bar in the lag plot indicates the relative lag between lLNvs and sLNvs, satisfying the experimental constraint. Schemes on the right illustrate corresponding observables.

We can use the theory to further investigate how the forcing affects observables (Fig. S7). We focus on detuning and forcing strength since these are key determinants of the forced oscillatory dynamics (Fig. 7B). Amplitude is not strongly affected by detuning, but grows steadily with forcing strength. The time lag increases with increasing detuning and is reduced with increasing forcing strength. The resulting relative time lag between lLNvs and sLNvs is approximately preserved for varying forcing frequency Ω and becomes smaller with increasing forcing strength. Finally, relaxation time is strongly controlled by forcing strength. Fast relaxation times require a large forcing strength, which results in a flat theoretical PRC (Fig. 5C), consistent with the experimental observation (Fig. 3B).

These results show that the generic theory is consistent with our experimental data for both lLNvs and sLNvs, showing quantitative agreement with measurements of amplitude, relaxation time, and relative lag between small and large LNvs.

### LNv oscillations are also synchronized with other neurons

Finally, we investigated whether synchronization of membrane oscillations was only circumscribed to the LNvs, or if it was a more widespread phenomenon. We performed simultaneous patch clamp recordings of pairs of neurons composed of one LNv and a non fluorescently labeled neuron localized nearby, in a distance range of approximately 15 to 30 *µ*m within the accessory medulla. We termed these neurons accessory medulla PDF-negative, aMe-PDF(-), because we did not have information about the precise nature of these cells other than their anatomical position and the fact that they did not express PDF.

We found coherent membrane oscillations between aMe-PDF(-) and LNvs, both with lLNvs (Fig. 8A, C) and with sLNvs (Fig. 8B, D), in all neuronal pairs that we were able to record from. Using cross-correlation analysis and the threshold crossing method, we consistently uncovered different lags between the LNvs and the aMe-PDF(-) cells. Membrane oscillations of lLNvs and aMe-PDF(-)s presented predominantly negative small lags (Fig. 8E). In the case of the sLNvs, four out of five aMe-PDF(-)s presented positive lags (were ahead of the sLNvs) and one aMe-PDF(-) had a negative lag (was delayed with respect to the sLNv) (Fig. 8F). These experiments suggest that membrane potential oscillation is a widespread phenomenon among aMe neurons.

**Figure 8.**
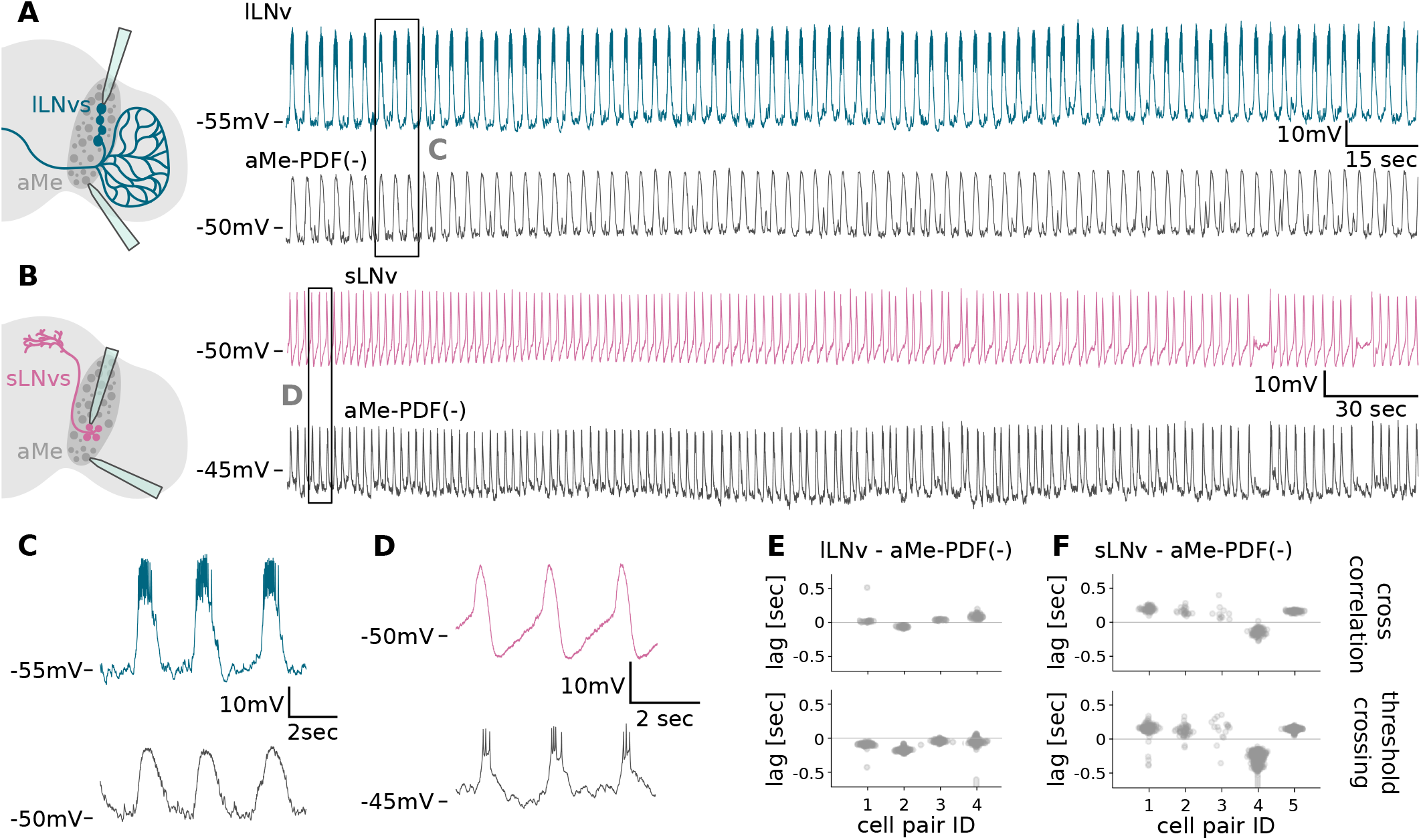
Other aMe neurons also oscillate coherently with LNvs. (A, B) Left: scheme of brain explant displaying aMe neurons together with (A) lLNvs and (B) sLNvs, with recording electrodes. Dark grey shade indicates the aMe region. Right: Fragment of representative dual recording from (top) LNv and (bottom) aMe-PDF(-) neuron, beginning (A) 32 min and (B) 28 min after dissection. Boxes mark zooms in C and D. (C, D) Representative fragments from boxes in A and B. (E, F) Lags obtained from (top) local cross correlation and (bottom) threshold crossing analysis for individual dual recordings of aMe-PDF(-) neurons and (E) lLNvs and (F) sLNvs. Recordings in A, B correspond to pair 4 in E and 5 in F.

## DISCUSSION

By means of simultaneous patch clamp recordings of pairs of neurons in the aMe, we reveal that this area of the *Drosophila* brain harbors neurons with coherent oscillation of their membrane potential. We focus on LNvs, a group of neurons with recognized roles in circadian timekeeping and sleep/wake behavior. We show that the membrane potential of lLNvs and sLNvs oscillates consistently during long term ex vivo recordings. lLNvs display a flat phase response curve, and both LNv types required the neurotransmitter ACh to sustain oscillatory activity, suggesting that oscillations are not cell-autonomous. Based on these data, we proposed a generic forced oscillator theory. The theory further predicts that neurons of the same type should oscillate in synchrony, and that a phase lag could occur between different neuronal types. We confirmed these predictions using dual recordings, observing that sLNvs oscillate behind lLNvs with a consistent lag. We then recorded cell pairs composed of an LNv and an unidentified aMe neuron, and revealed that they were also coordinated, with varied lags. This observation motivates the hypothesis of collective rhythms in the aMe, resulting from the synchronization of the membrane oscillations of local neurons. Further work is required to test this hypothesis, and to unravel the function of this putative emergent property of the aMe.

### The source of collective neuronal oscillations

Our results provide strong evidence for synchronized membrane potential oscillation of clock neurons in *Drosophila*, in line with a recent report [19]. Moreover, several lines of evidence indicate that the frequency of these oscillations depends on intact visual circuits. First, the oscillation frequency of LNvs decays in ex vivo preparations (Figs. 1 and 2) [18], presumably due to disruption of upstream visual input circuits. Second, this frequency decay is slower in ex vivo semi-intact preparations [18], where visual inputs are relatively preserved. And third, oscillation frequency is higher in LNvs recorded in vivo [19], where visual inputs are not altered. Additionally, our quantification of frequency slowing down carries practical implications for ex vivo measurements generally. We argue that it is important to document the time since dissection in assays performed on brain explants. To quantitatively compare the effects of different conditions or genotypes on physiology, it is essential to obtain recordings at roughly the same time since dissection.

Several observations support the idea that synchronized oscillations of clock neurons originate from shared cholinergic inputs (Figs. 3 and 4) [18, 19]. ACh is the most prevalent neurotransmitter in fly visual processing circuits [46]. A potential origin of ACh could be a subset of aMe neurons labeled by VT037867-Gal4 driver, that have been described to be presynaptic partners of clock neurons [19, 50]. Whether these cells are also providing cholinergic input to other, non-circadian, aMe neurons should also be determined in the future.

### A role for phase lags in traveling waves

We found an overall coherent oscillation in all neuronal pairs recorded, showing coordinated membrane potential cycles throughout the recordings. When recording from two neurons belonging to the same LNv type we found strong synchronization with zero lag. In contrast, in recordings from different neuronal type pairs we observed consistent time lags, with sLNvs always lagging 0.10 to 0.16 sec behind lLNvs (Fig. 6). While in our hands the lag between lLNvs and sLNvs oscillations is apparent in voltage traces, and consistently quantified in multiple ways, a previous work did not report such time delays in clock neuron oscillations [19]. We also found lags between LNvs and other neurons from the aMe, suggesting that this may be a widespread feature of the network.

Since the lag between lLNvs and sLNvs is in the order of 100 milliseconds, we expect a lag with the driver to be at least in the same order of magnitude. What could be causing such lag with the driver? Synaptic connectivity of the whole *Drosophila* brain has recently been revealed through a connectomics approach [51]. However, the lag timescale is hard to reconcile with individual classical chemical synapses, which are about two orders of magnitude faster.

Another possibility is that the lag is caused by distinct cholinergic ways of communication. While ACh neurotransmission can be very fast, like at the mammalian neuromuscular junction, it can also be slower, depending on the type of ACh receptor expressed in the postsynaptic neuron, the location of receptors relative to the ACh release site, and the concentration of acetylcholinesterase in the extracellular space [52]. At least ten nicotinic ACh receptor genes have been reported in *Drosophila*, which encode different subunits that can be combined to produce the mature pentameric ACh receptors [53]. Thus, differences in cholinergic input processing could cause differing lags in different neuronal types.

In addition to cholinergic inputs, the lag may involve peptidergic transmission that provide essential modulation to neuronal physiology. Peptidergic neuromodulation is slower due to its volume transmission nature. Neuropeptides seem to be pervasive in the insect aMe [54, 55], and many neuropeptide receptors are expressed in clock neurons [56]. However, due to their volumetric way of passing on information and the lack of versatile genetic tools, neuropeptides roles in neuronal physiology modulation within intact circuits is still under investigation.

While these hypotheses deserve further exploration, here we proposed a theoretical description that captures aspects of membrane potential oscillations. In the framework of this theory, lags naturally arise between each neuron and an external forcing. These lags are determined by parameters that encompass physiological properties of neurons. For example, membrane time constant could influence the characteristic frequency scale *ω*, and neurotransmitter signal processing and their modulation might affect the forcing strength *f* in the theory. Since different neuronal types may differ in these physiological properties, we expect them to display diverse lags with respect to the forcing. Additionally, the theory makes predictions that could be tested experimentally, such as the effects of decreasing forcing strength on relative lags and oscillation amplitude. In summary, the theory provides a parsimonious explanation for the lag we observe between pairs of aMe neurons, arising from the different biophysical properties of the neurons involved.

If there were no lags between pairs of neurons, we would expect a collective rhythm in which all neuronal types oscillate in synchrony. The presence of consistent phase lags between groups of oscillators, as we report here, hints at the possibility that traveling waves of activity may be present in the fly brain. Brain waves may be relevant to time sequences of neuronal activation, something that has been argued to be crucial for brain function [10, 11, 57]. Several mechanisms have been proposed to account for wave propagation in the context of neuronal circuits, such as (i) locally coupled autonomous oscillators, (ii) propagating pulses from a single driving oscillator through a sequence of synapses, and (iii) a single oscillator directly driving target neurons with increasing delays [57]. Here we propose the additional possibility, (iv) that neuronal biophysical properties determine the phase lag for each target neuron with respect to an oscillating driver. Wave propagation could occur across three dimensional space, or following the more complex neuronal network topology. Local field potential recordings using multielectrodes, suggested that oscillations may travel from one brain location to another, providing indirect evidence for waves in the fly brain [15]. A more direct evidence of brain waves could be obtained by high resolution mesoscopic imaging using voltage sensors. Mapping phase lags in other oscillating components of the fly brain would also provide strong evidence for traveling waves.

### Are oscillations in the *Drosophila* aMe part of a larger collective phenomenon?

The aMe is a strategically situated brain area, as it is located in a constriction of the tissue between the optic lobes and the central brain. Similar to many other animals, flies rely heavily on vision to sense their environment. In *Drosophila*, more than half of the brain’s neurons are on the optic lobes [58], where they organize in stratified neuropiles devoted to visual processing. Visual information has to inescapably traverse the aMe to reach the central brain, where it is integrated with other sensory modalities and stored information, to ultimately induce motor behaviors [59]. Axonal projection pathways from visual processing neurons that go through the aMe and reach the central brain have been described. For instance, visual features are passed on to optic glomeruli of the posterior lateral protocerebrum; visual context is relayed to the mushroom bodies for visual memory formation; and visuo-spatial information is conveyed to the central complex, to aid navigation [46]. But the aMe region does not only host axonal projections, many neuronal somas are located there as well. Out of the many aMe neurons, the only ones with characterized functions, to the best of our knowledge, are the few ones belonging to circadian circuits. The roles of most other aMe resident neurons have not been revealed yet, but we here describe that many of them present synchronized oscillations of their membrane potential. Additional efforts should delve into the functions of aMe neurons besides circadian control, and into the meaning of their coherent oscillation.

It is possible that the predicted aMe waves could extend to other regions of the *Drosophila* brain. This is suggested by different lines of evidence, for example clock neuron clusters localized outside the aMe in dorsal brain regions (DN1a, DN3a, DN3p) were shown to oscillate with a similar frequency (0.3-0.4Hz) [19]. While synchrony between these dorsal clock neurons and aMe neurons has not been examined, the consistent oscillation frequency suggests a potential connection. Neuronal synchronization has also been reported in the sleep-regulating R5 neurons [16], with an oscillatory frequency in the same range to the one we described. Overall, evidence indicates that a number of neurons in the *Drosophila* brain are oscillatory in nature and present membrane potential synchronization.

### Is the aMe the second hand of the *Drosophila* clock?

Neuronal oscillation provides an extra layer of complexity for neuronal information integration [10]. This occurs because the impact of incoming information, conveyed through the arrival of neurotransmitters released by upstream neurons, may vary based on the phase of the oscillatory cycle when the synaptic input is received. Membrane potential has been shown to modulate ionotropic [60, 61] and metabotropic receptors [62, 63], and it determines the driving force of the ions that permeate through ligand-gated ion channels. In a non-oscillating neuron, its output is solely determined by the spatial and temporal summation of received inputs. Conversely, in oscillatory neurons, the impact of identical inputs may differ based on the phase of the cycle when the neuron receives them.

LNvs are highly integrative neurons. Their role in integrating information is illustrated by the multitude of information sources they receive: not only from the visual circuits [18, 50], but also from extra-retinal photoreceptors [64], from light itself [65–68] and via synaptic inputs from numerous clock and non clock neurons [69]. Their oscillatory activity should be kept in mind when modeling the integration of all these information sources. For decades, it has been acknowledged that the *Drosophila* aMe contains crucial components of the 24-hour clock, a timeframe aligned with the Earth’s rotation. Predicting events at shorter timescales has advantages too; however, the cellular, molecular and circuital mechanisms of ultradian clocks remain less explored [70, 71]. The estimation of time in the range of seconds has recently been described in *Drosophila* [72]. Could the membrane oscillations of aMe neurons be part of the second hand of the fly clock? Only time will tell.

## MATERIALS AND METHODS

### Experimental model and subject details

*Drosophila melanogaster* flies were grown and maintained at 25^°^C in standard cornmeal medium under 12/12 h light/dark cycles. 3-to-9 day old female Pdf-RFP flies [39] were used for all the experiments.

### Electrophysiology

Patch clamp experiments were conducted as previously described [17, 36–38]. Briefly, 3-to-9 day old female flies were anesthetized with a short incubation of the vial on ice, and brain dissection was performed in external recording solution, which consisted of the following, in mM: 101 NaCl, 3 KCl, 1 CaCl2, 4 MgCl2, 1.25 NaH2PO4, 5 glucose, and 20.7 NaHCO3, pH 7.2, with an osmolarity of 250 mmol/kg. After removal of the proboscis, air sacks, and head cuticle, the brain was glued ventral side up to a Sylgard-coated coverslip using a few microliters of tissue adhesive Vetbond (3M). The time from anesthesia to the establishment of the recordings is referred as “time since dissection”, which was spent as follows: LNvs were visualized by red fluorescence in Pdf-RFP flies, which express a red fluorophore under the Pdf promoter [39], using an Olympus BX51WI upright microscope with 60× water-immersion lens, and ThorLabs LEDD1B and TK-LED (TOLKET S.R.L.) illumination systems. Once the fluorescent cells were identified, cells were visualized under IR-DIC using a DMK23UP1300 Imaging Source camera and IC Capture 2.4 software. lLNvs were distinguished from sLNvs by their size and anatomic position. To allow access of the recording electrode, the superficial glia directly adjacent to LNvs somas were locally digested with protease XIV solution (10 mg/ml; P5147, Sigma-Aldrich) dissolved in external recording solution. This was achieved using a large opened tip (∼20 *µ*m) glass capillary (pulled from glass of the type FG-GBF150-110–7.5; Sutter Instrument) and gentle massage of the superficial glia with mouth suction to render the underlying cell bodies accessible for the recording electrode with minimum disruption of the neuronal circuits. After this procedure, the protease solution was quickly washed by perfusion of the external solution using a peristaltic pump (catalog #ISM831, ISMATEC). Recordings were performed using thick-walled borosilicate glass pipettes (FG-GBF150-86-7.5, Sutter Instrument) pulled to 7-8 MΩ using a horizontal puller P-97 (Sutter Instrument) and fire polished to 9-12 MΩ. Recordings were made using a Multiclamp 700B amplifier controlled by pClamp 10.4 software via an Axon Digidata 1515 analog-to-digital converter (Molecular Devices) and saved into .abf files. Recording pipettes were filled with internal solution containing the following (in mM): 102 potassium gluconate, 17 NaCl, 0.085 CaCl2, 0.94 EGTA, and 8.5 HEPES, pH 7.2, with an osmolarity of 235 mmol/kg. Gigaohm seals were accomplished using minimal suction followed by break-in into the whole-cell configuration using gentle suction in voltage-clamp mode with a holding voltage of −60 mV. Spontaneous firing was recorded in current clamp (*I* = 0) mode. For the dual recording experiments, the sequence was done two consecutive times, achieving the whole-cell patch clamp configuration in one neuron after the other (in the case of sLNv-lLNv recordings, the lLNv was patched in the first place). Once both neurons were patched, the electrical activity was recorded for both neurons simultaneously. Simultaneous recordings were always performed in neurons located ipsilaterally. For the mecamylamine experiments, after 1 to 3 min of recording basal conditions, 10 ml of mecamylamine (M9020, Sigma-Aldrich) solution 10 *µ*M prepared in external saline was perfused over ∼3 min. Mecamylamine was then washed out with external saline perfusion.

For the phase response curve experiments, neurons were initially recorded for 1 to 3 minutes to determine their baseline bursting frequency (bursts per 60 seconds). The stimulation frequency was then calculated by dividing the baseline bursting frequency by 5. To ensure that the stimulation frequency did not coincide with the neuron’s spontaneous firing, the value was rounded, which resulted in a shift of ± 0.01 to 0.02 Hz. This adjustment prevented the stimulation frequency from being an exact multiple of the baseline firing frequency, so that stimulation would be unlikely to fall in the same part of the cycle repeatedly. Once the stimulation frequency was determined, neuronal stimulation was performed using pulses of -20 pA amplitude and 100 ms duration. Each stimulation trial consisted of 5 sweeps, each lasting 90 seconds. After the final sweep of stimulation, it was verified that the neuron continued to fire spontaneous bursts with a frequency similar to the baseline frequency recorded at the beginning.

### Code

The analysis of electrophysiological recordings was performed using custom python code with the standard scientific libraries (scipy, numpy, pandas, matplotlib) and the pyABF [73] library to read the .abf files, and the jitcode library for numerical integration [74].

### Data filtering

Non-causal Butterworth filters of order 2 were applied as implemented in scipy.signal.filtfilt, to detrend and denoise the data (Fig. S1A). Detrending was performed using a highpass filter with cutoff frequency 0.1Hz (Fig. S1B), termed filt1. Denoising was done with a lowpass filter with cutoff frequency 10Hz, termed filt2. This also results in the loss of action potential spikes (Fig. S1C). The peak finding step implemented for some calculations below required additional smoothing, which we did using a second lowpass filter with frequency cutoff at 2Hz (Fig. S1D), termed filt3.

### Threshold crossings and cycle duration

We developed an approach to reliably detect threshold crossings in the presence of fluctuations and variability observed in the data. We located cycle peaks in the detrended and smoothed data (filt1, filt2 and filt3), by first finding all local maxima with `scipy.signal.find_peaks`, and then filtering the ones that had a prominence above 3mV, and that were at least 0.4 times the average distance between local maxima (Fig. S1E). The former prevents shallow, spurious peaks in between pulses from being detected, and the latter prevents more than one peak per cycle from being detected.

Fluctuations in the shape of the cycle cause fluctuations in the timing of the cycle peak. Thus, distance between peaks is not a good estimate for cycle duration (CD). For this reason, we detected rising edge threshold crossings in the detrended and denoised data (filt1 and filt2). Starting at the position of each peak in the smoothed data, we backtrack until we find a point that crosses a given threshold. To avoid arbitrariness in defining the threshold, we set 21 threshold values between 0mV and 10mV in the detrended signal (Fig. 1E, Fig. 2C), which encompasses most of the signal range for most recordings. CD values were calculated as the difference between threshold crossing times of two consecutive cycles, for each threshold value. Mean and SD were then calculated from this set of CD values.

### Valid cycle filtering

For each recording, individual CDs can be plotted as a function of time. While CD steadily increases in most cases, a few points deviate systematically from the global trend, with values that are about twice those surrounding them (Fig. 1C, E). Inspecting the data in the vicinity of these points reveals that they mostly occur when either the neuron appears to skip a cycle (Fig. S5), or when the cycle amplitude was relatively small and the algorithm failed to detect it (Fig. 1C). These non valid cycles are in some cases interesting behaviors, but they are not representative of the global trend. To filter them out we use a two step method. First, we calculate the population mode of the CD, and discard points that are above 1.8 times the mode. However, this step alone is prone to discarding points that are valid cycles if the recording is too long, due to the CD drift. Performing this test locally for shorter windows of recordings is not a good alternative, since this step relies on most cycles being valid to have a good estimate of the mode and non valid cycles are usually closer to the end of the recording (Fig. S2). In the second step, we calculate the trend in the remaining points by linear fitting. Then, we subtract the trend from the whole data set, including the previously discarded points, and add the value of the trend at zero. Then the mode is calculated again, and points above 1.8 times the mode are considered non valid cycles. With this method, around 97% of the data is considered valid around the beginning of each recording on average, dropping to about 93% at the end of the recordings (Fig. S2C, D).

### Initial cycle duration

Initial cycle duration of a recording is calculated by averaging the first 40 valid cycles.

### Baseline

To calculate the baseline of a recording, we first denoised the signal (filt2). We located local minima using scipy.signal.find peaks, and discarded those above the mode. With this approach, the remaining local minima correspond mostly to fluctuations around the membrane potential baseline. We applied this protocol in 20 sec non-overlapping windows to determine a local value of the membrane potential baseline (Fig. S3A).

### Amplitude

We determined an amplitude value for each individual cycle in a recording. We defined the amplitude as the distance from the peak of the cycle to the local baseline (Fig. S4A). Since the shape of the cycle does not strongly affect the amplitude, smoothing was applied (filt3) to identify cycle peaks, but not for the baseline calculation.

### Phase Response Curve

To obtain a phase response curve from current injection experiments, we first processed recordings following the protocol to find threshold crossings above. Here we defined a cycle as the time interval between two consecutive threshold crossings, and selected cycles with a stimulus. We calculated the ratios

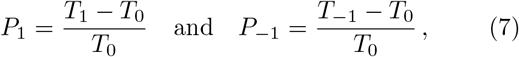

where *T*_1_ is the CD of a cycle where the stimulus was applied, *T*_0_ the previous one and *T*_*−*1_ the one before that. The ratio *P*_1_ is a phase shift due to the stimulus, which is positive if the perturbed cycle is longer than the basal cycle, and negative if it is shorter (color dots in Fig. 3B). The *T*_0_ in the denominator normalizes the differences between consecutive cycles, to account for the increase in CD as a function of time since dissection. To control for intrinsic fluctuations in CD, we compare the value of *P*_1_ to *P*_*−*1_, which represents the basal variability in the absence of a perturbation with respect to the reference cycle *T*_0_.

To determine the phase of the cycle at which the perturbation was applied, we map the duration of the cycle to the interval [0, 2*π*], and the timing of the stimulus start is linearly mapped in that interval [45]. The shaded area displays the standard deviation of all *P*_*−*1_ values, representing a detection limit for the measurements.

### Relaxation time

We identified cycles where perturbations were performed as above. To calculate relaxation times, we selected cycles where the phase of the perturbation was between 0.6*π* and 1.6*π*. This ensures that the perturbation happens during the flatter part of the cycle, where the impulse response is cleaner. We fitted the relaxation with an exponential function, and obtained the relaxation time *t*_*r*_ as the characteristic timescale of the exponential decay (Fig. 5D).

### Pulsing rate

To calculate the pulsing rate, we detected cycles using the protocol for threshold crossings. We then counted the number of cycles in 60 second windows, with 30% overlap between contiguous windows.

### Lag

To obtain cross correlation lags from detrended and denoised data (filt1 and filt2), we first calculate the cross correlation *C* between two channels *u*(*t*) and *v*(*t*),

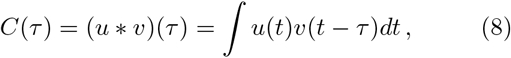

where ∗indicates a convolution and the integral runs over the recording time, and normalize by the maximum value of *C*(*τ*). We implement this using the correlate and correlation lags functions in the scipy.signal library. We defined the lag between channels in dual recordings as the correlation lag at maximum correlation. We determined a local lag value from the cross correlation analysis applied on 10 sec non-overlapping windows.

To obtain threshold crossing lags, we first determined threshold crossings in both channels of the dual recording following the threshold crossings protocol above. Then, for each crossing in one channel, we searched for the closest crossing in the other. This selection is not ambiguous because the protocol ensures that only one crossing per cycle is selected. Since in some cases a pulse is not detected, we discarded matching cycle pairs that are separated by more than 0.4 times the local cycle duration. This amounts to pulses between 0.8 and 1.4 sec apart, depending on the time since dissection, values which are significantly larger than every lag we measured.

## SUPPLEMENTAL METHODS

### Forced oscillator

Here we formulate the generic theory for forced oscillations. The loss of membrane potential oscillations after mecamylamine (mec) application, blocking acetylcholine receptors, suggest that the oscillations are driven by an external source. In absence of the driver, the oscillations stop. Furthermore, the flat PRC could be explained by a strong coupling with the driver, such that after perturbation the oscillator phase in quickly re-entrained. Many neuronal models describing ion channels can give rise to oscillations through a supercritical Hopf bifurcation [43]. Thus, here we choose a generic model with a Hopf bifurcation and forcing,

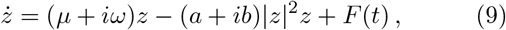

where *z* is a complex variable, *µ* is the bifurcation parameter, *ω* a characteristic frequency, *a* controls the convergence to the limit cycle, and *b* is an amplitude dependent correction to the frequency termed non-isochronicity of the oscillator. The forcing is

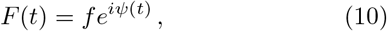

with forcing strength *f* controlling the coupling to the driver, and forcing phase

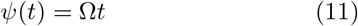

growing linearly with speed Ω. Here we are interested in the case *µ <* 0, such that in the absence of forcing, *f* = 0, there are no autonomous oscillations. We also require *a, ω*, Ω, *f >* 0.

Writing *z* = *re*^*iθ*(*t*)^, the polar form of Eq. (9) is

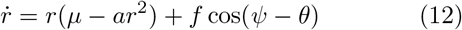

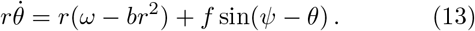

Stepping into a rotating reference frame, we introduce a phase relative to the driver *ϕ* = *ψ − θ*, such that

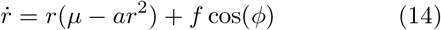

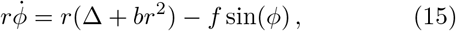

where we defined the detuning Δ *≡* Ω − *ω* between driver and oscillator.

### Equilibrium

Next, we look for the equilibrium given by

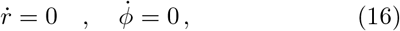

such that the oscillator has a constant limit cycle radius *r*(*t*) = *r*_0_ and phase *ϕ*(*t*) = *δ*. This means that in equilibrium, the instantaneous frequency in the original reference frame is 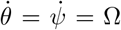, representing the entrainment of the oscillation to the forcing with a constant lag between forcing and oscillator

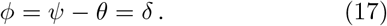

Replacing Eq. (16) into Eqs. (14) and (15),

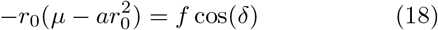

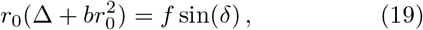

we obtain the expressions for the equilibrium variables

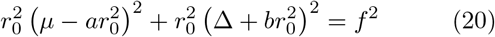

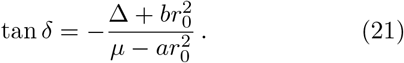

Eq. (20) can be cast into a cubic equation in terms of 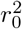, and solved analytically by means of Cardano’s formula to show that for *µ <* 0 there is always one real positive root.

### Linear stability analysis

Next, we perform a linear stability analysis of the equilibrium state, to characterize the response of the system to perturbations. We consider perturbations around the equilibrium *dr*(*t*) ≡ *r*(*t*) − *r*_0_ and *dϕ*(*t*) ≡ *ϕ*(*t*) −*δ*. For small perturbations, we discard higher order terms and obtain the linearized system

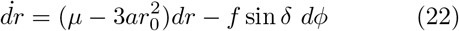

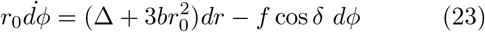

with Jacobian matrix

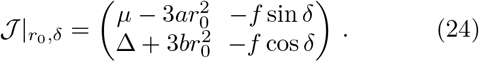

Using Eqs. (18) and (19) we can write the trace and determinant of the Jacobian matrix as

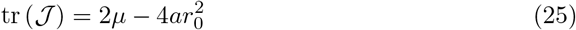

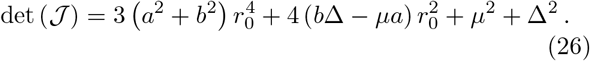

Since tr (*J*) *<* 0 and det (*J*) *>* 0, the equilibrium will always be stable [49]. Eigenvalues *λ*_*±*_ can be obtained as

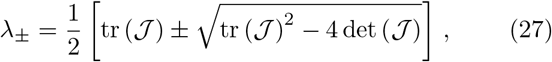

which results in

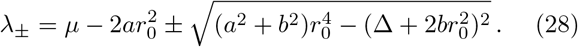

These eigenvalues describe the relaxation back to equilibrium along the directions given by the corresponding eigenvectors

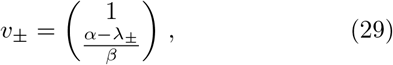

with

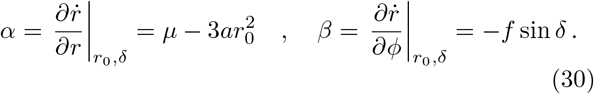

### Perturbation response

To obtain the experimental PRC, we inject current to the neuron. This perturbs the equilibrium value of the membrane potential, which quickly returns to steady state cycling. It is convenient to interpret one of the Cartesian coordinates

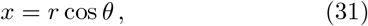

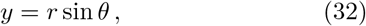

of the complex variable *z* = *x* + *iy* as this membrane potential. Without loss of generality, here we assume the membrane potential is described by Eq. (32), so a negative current injection results in a negative perturbation in *y*. Assuming the system is in equilibrium before a perturbation at time *t*_0_, we can write the perturbed state (*r*_*p*_, *θ*_*p*_) as

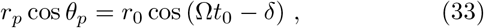

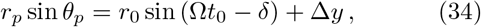

where Δ*y <* 0 is the instantaneous perturbation. The perturbed state at *t*_0_ satisfies

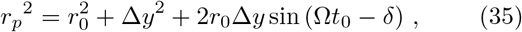

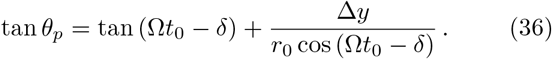

The time evolution of this perturbation can be written in terms of eigenvectors and eigenvalues as

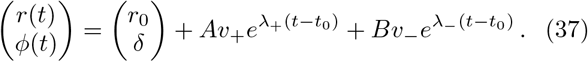

Since |Re(*λ*_+_)| *<* |Re(*λ*_*−*_)|, long term perturbation dynamics is dominated by the slower timescale given by | Re(*λ*_+_)|^*−*1^. Thus, we approximate

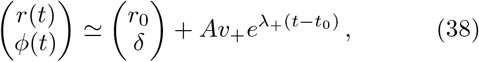

and associate the experimental relaxation time to this eigenvalue through the expression

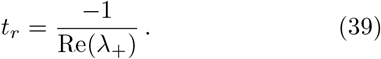

### Parameter values

Next, we look for parameter values that can account for lLNvs and sLNvs data. We assume that the forcing frequency Ω in Eq. (11) is the same for different neuronal types, based on the observation of coherent oscillations. Assuming that the recordings are in steady state, so oscillations are entrained to the forcing, Eq. (17), we determine the frequency Ω from CD measurements as

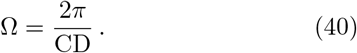

Since the value of CD changes over time, we choose a time point 25 min after dissection for our measurements, resulting in CD = 2.20 sec (Figs. 1 and 2 of the main text). The polar form of the forced oscillator, Eqs. (12) and (13), includes five parameters that we need to determine for each neuronal type,

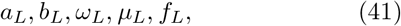

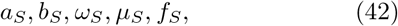

where we label *L* and *S* parameters that correspond to lLNvs and sLNvs, respectively. Unlike the forcing frequency, forcing strength *f* may differ for lLNvs and sLNvs, since they may sense the signal differently. In this work, we set *b*_*L*_ = *b*_*S*_ = 0 for simplicity.

We can relate parameter values to observables of the theory, the amplitudes *r*_0_, time lags *τ* = *δ/*Ω and relaxation times *t*_*r*_ = *−*1*/λ*_+_ for lLNvs and sLNvs,

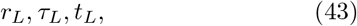

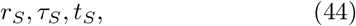

through equillibrium solutions Eqs. (18) and (19), and linear stability Eq. (28) for both neuronal types. Between parameters and observables, Eqs. (41) to (44), we have 16 unknowns. We obtain the equillibrium amplitude *r*_0_ from the measured membrane potential amplitude 2*r*_0_ for both neuronal types (Fig. S4C), *r*_*L*_ = 11.4mV and *r*_*S*_ = 8.9mV. Although we do not have direct measurements of time lags with the driver, we did measure the relative time lag between sLNvs and lLNvs, *τ*_*S*_ *− τ*_*L*_ = 0.13 sec (Fig. 6 of the main text). We measured the relaxation time for lLNvs *t*_*L*_ = 0.023 CD (Fig. 5E of the main text). Since we did not measure the relaxation time in sLNvs, we assume *t*_*S*_ = *t*_*L*_ for simplicity.

After measurements and assumptions, 9 unknowns remain. Applying Eqs. (18), (19) and (28) to both neuronal types provides 6 additional constraints, leaving 3 degrees of freedom among parameters and remaining observables. This means that to complete the parametrization of the theory, we need to choose values for 3 of the unknowns. For some of these unknowns we can make educated guesses. For example, we assume characteristic frequencies *ω*_*L*_ and *ω*_*S*_ to be similar to the forcing frequency Ω. With this assumption, we can expect the phase lags to be small relative to the cycle, Eq. (21). Additionally, measurements of relative lags to other neuronal types produces consistently bounded values (Fig. 8). Setting the value of *δ*_*L*_, *δ*_*S*_ is constrained by the measured relative lag.

We invert Eqs. (18), (19) and (28) to obtain expressions for the remaining parameters

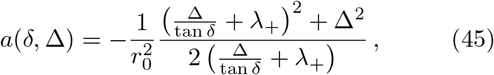

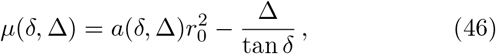

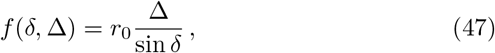

where we omitted labels *L* and *S* for readability. With these expressions, we explored how *a, µ* and *f* depend on *δ* and Δ for both neuronal types (Fig. 7A of the main text). A possible choice is that lLNvs are faster and ahead of the forcing, while sLNvs are slower and behind the forcing. For the remaining parameters, this choice results in values that are very similar between the two LNv types (Tab. I).

## ACKNOWLEDGEMENTS

We are grateful to Koichiro Uriu, Richard Baines and members of the Muraro Lab for critical reading of this manuscript. FFC and MW were funded by doctoral scholarships from CONICET. This work was supported by grants from AGENCIA I+D+i - FONCYT (PICT 2018-2030 and PICT 2020-1645 to NIM and PICT 2019-0445 to LGM), from CONICET (PIP 2022-2024 GI 11220210100779 CO to NIM), from the Chan Zuckerberg Initiative (CP2-1-0000000310 to NIM) and by FOCEM-Mercosur (COF 03/11) to IBioBA. The authors declare no competing financial interests.

## SUPPLEMENTAL FIGURES

**Figure S1.**
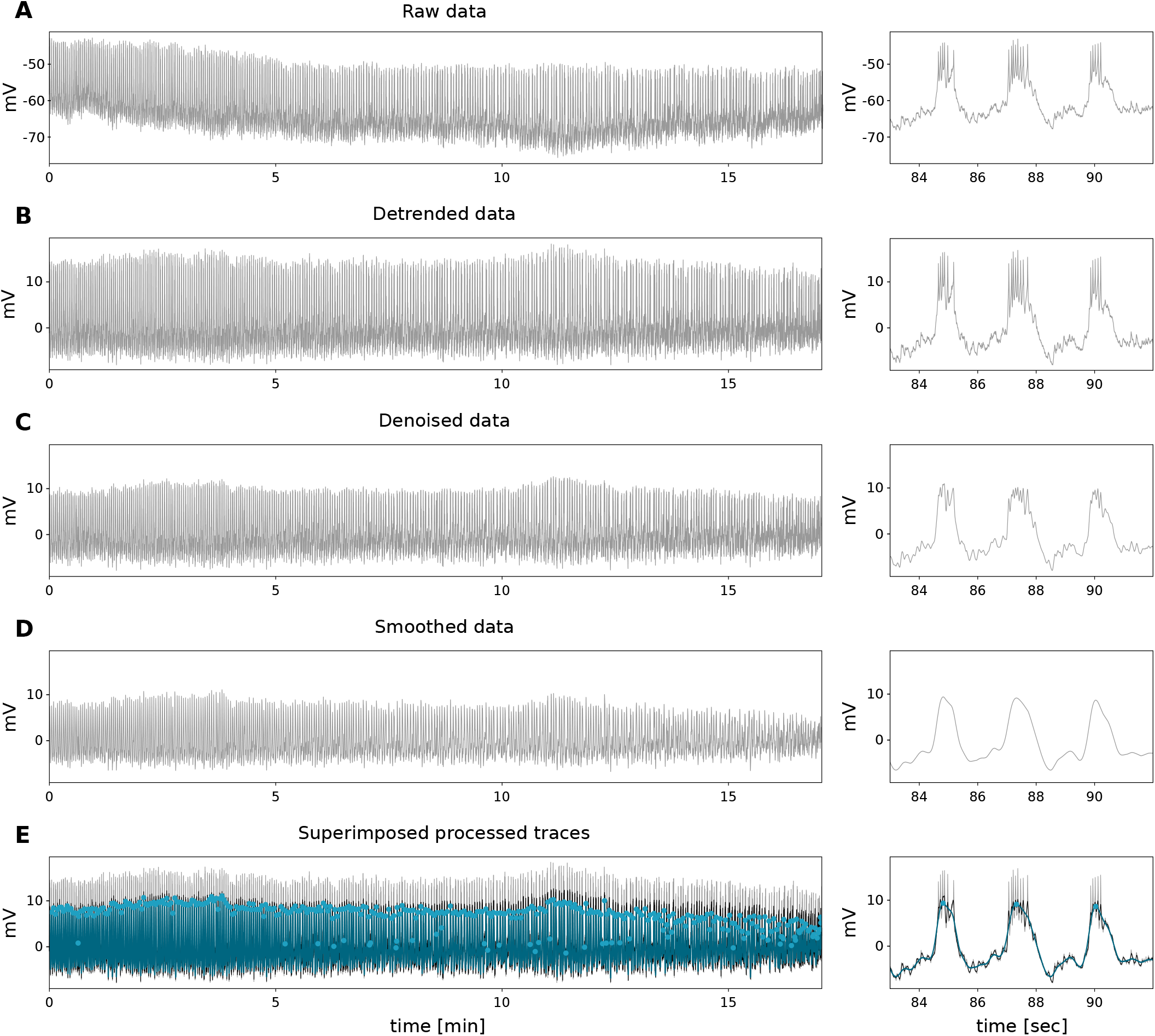
Processing of electrophysiological recordings. (A-E) Left: typical full recording. Right: detail. (A) Raw data. (B) Detrended data by highpass filter with cutoff frequency 0.1 Hz. (C) Denoised data by lowpass filter with cutoff frequency 10 Hz. (D) Smoothed data by a second lowpass filter with cutoff frequency 2 Hz. (E) Traces from B (grey), C (black) and D (blue) together with peak locations (dots) obtained from D. All filters are Butterworth filters of order two applied in forwards and backwards pass to avoid phase shifting.

**Figure S2.**
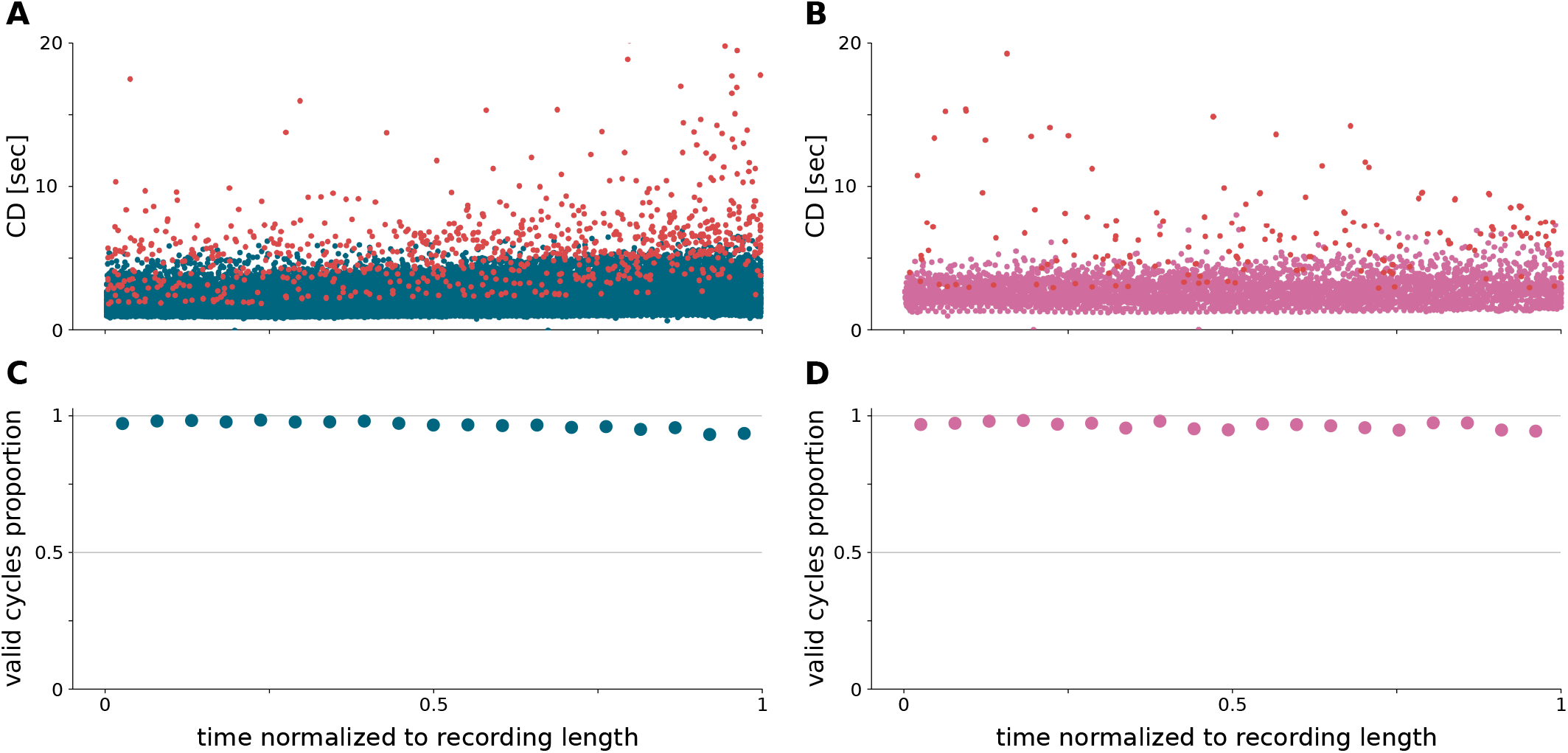
Non valid CDs represent a small fraction of the total. (A, B) Valid CD values from all (A) lLNv (blue dots) and (B) sLNv (pink dots) recordings, together with non valid CD values (red dots). Recording time was normalized to the total duration of each recording. (C, D) Fraction of valid points in windows of 5% of the total recording duration for (C) lLNvs and (D) sLNvs. The total fraction of valid CDs is always above 93%, averaging 96.7% of 25811 cycles for lLNvs and 96.4% of 5188 cycles for sLNvs.

**Figure S3.**
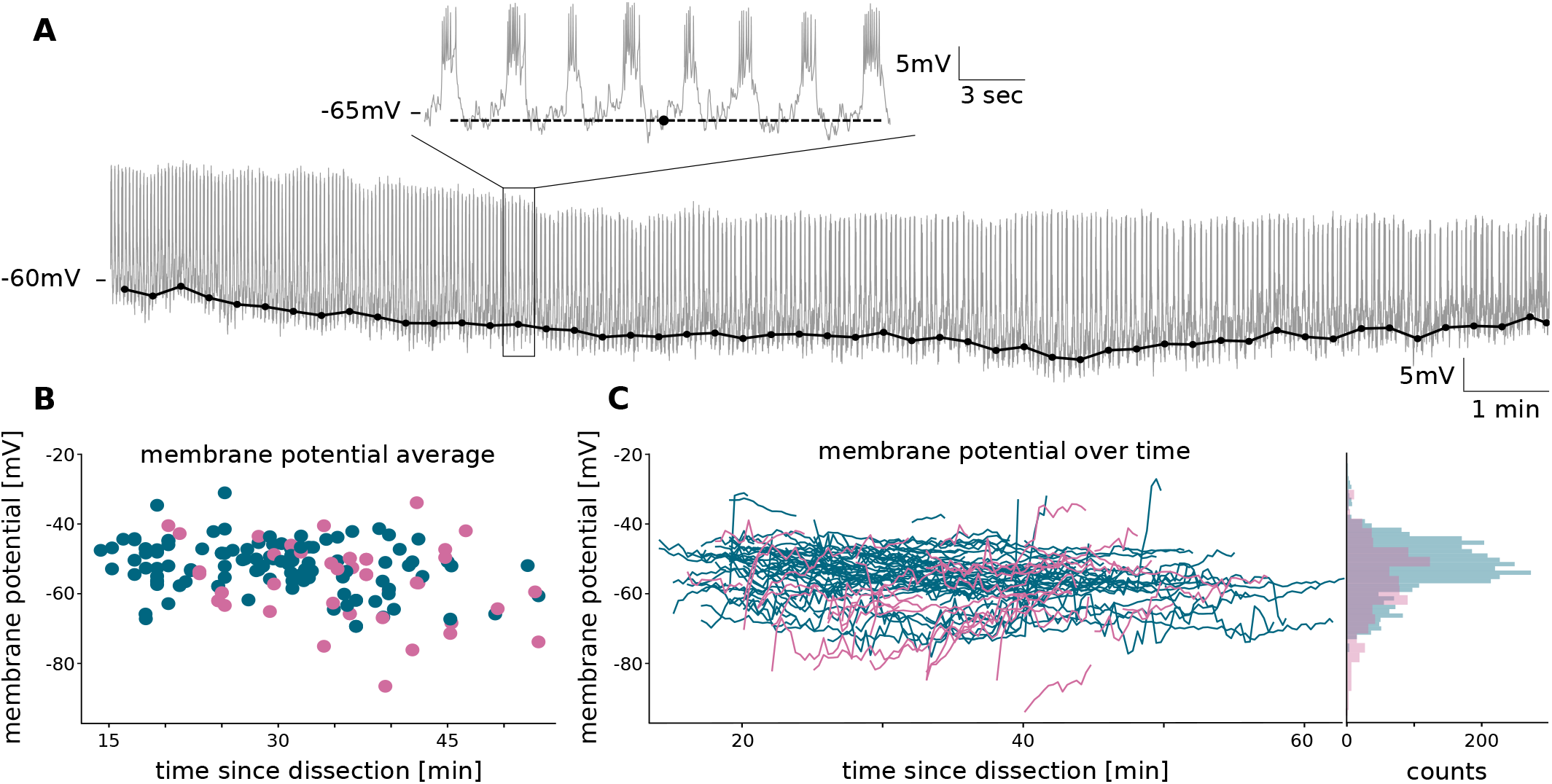
Baseline does not display a trend at the population level. (A) Representative example of membrane potential baseline calculation (black dots with joining lines) from raw data (grey line). Detail shows a 20 sec interval with the calculated baseline value (black dot) and horizontal dashed line for reference. (B) Average baseline value for all lLNv (blue dots) and sLNv (pink dots) recordings. The time coordinate for each point was chosen at the middle of the recording time. (C) Left: Baseline traces as a function of time since dissection from all lLNv (blue) and sLNv (pink) recordings. Right: Marginal distribution of baseline values from left panel.

**Figure S4.**
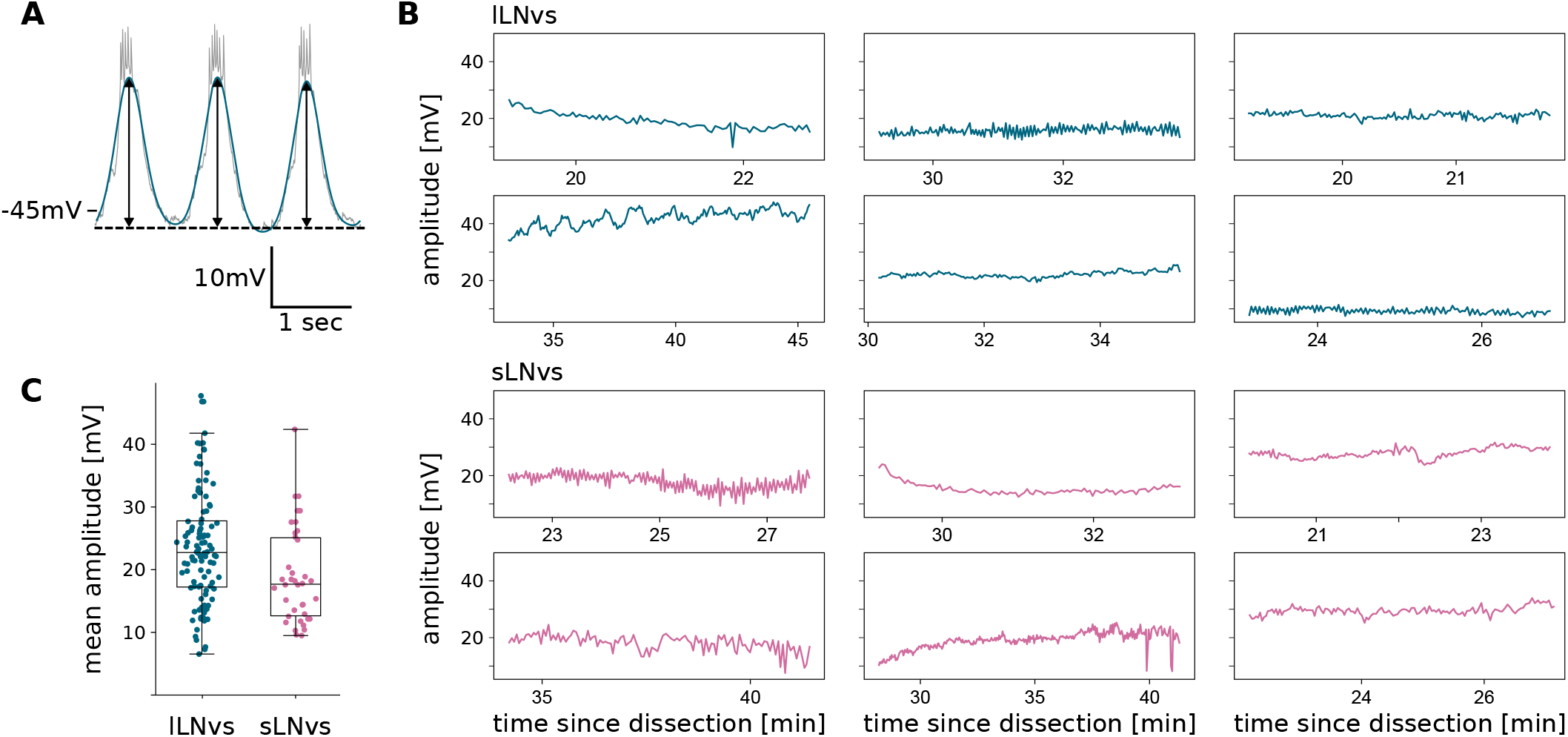
Amplitude does not display a trend at the population level. (A) Illustration of amplitude calculation from each cycle of membrane potential oscillations (grey line), obtained as distance (black arrows) from local baseline (dashed line) to the peak of the smoothed trace (blue line). (B) Representative examples of oscillation amplitude as a function of time since dissection for (top) lLNvs (blue lines) and (bottom) sLNvs (pink lines). (C) Mean amplitude value for all individual recordings from lLNvs (blue dots) and sLNvs (pink dots). Boxes are the interquartile range, bar is the median and whiskers extend to 1.5 times the interquartile range.

**Figure S5.**
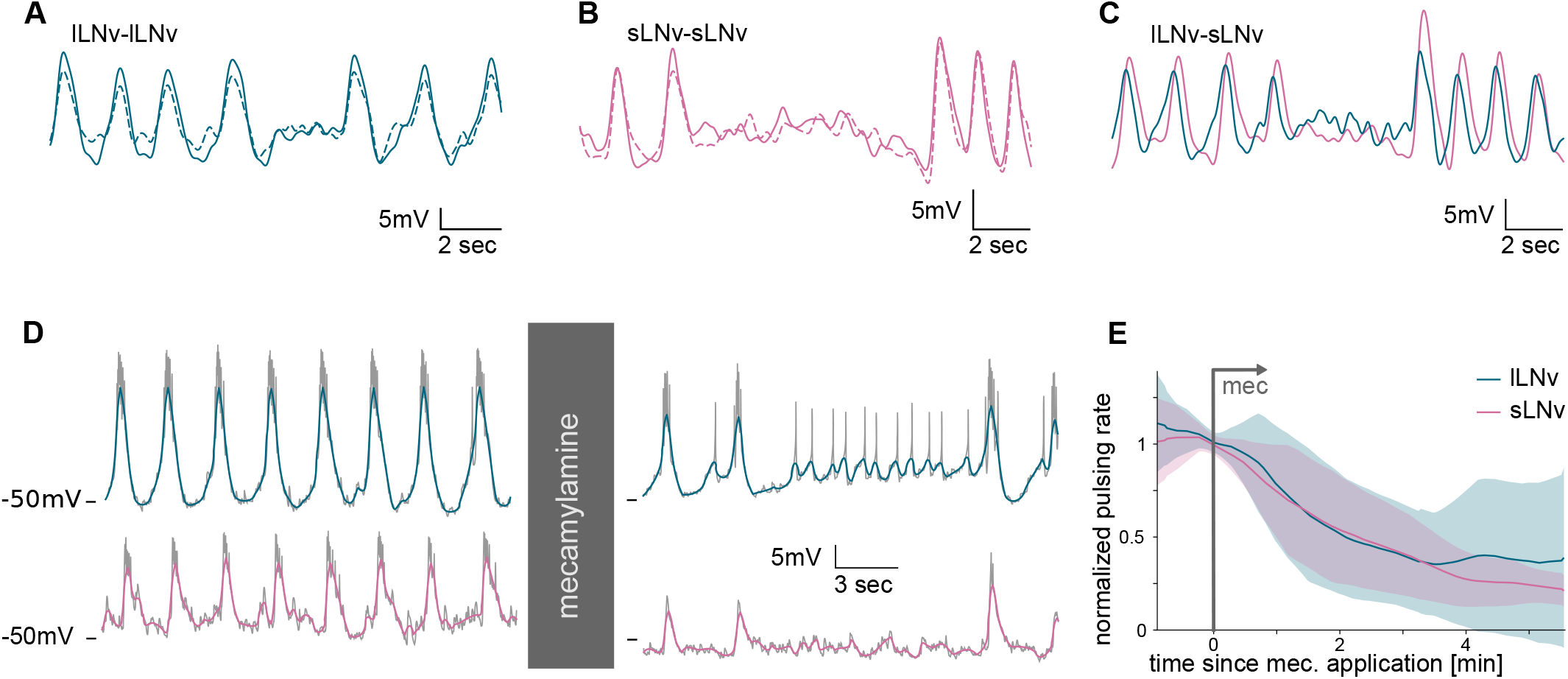
LNv oscillations are coordinated even through disruptions. (A-C) Representative detrended, denoised and smoothed dual recording fragments from (A) lLNv pair, (B) sLNv pair and (C) lLNv-sLNv pair. (D) Representative dual recording fragment (grey lines) and denoised traces (color lines) from a pair of (top) lLNv and (bottom) sLNv (left) before and (right) 120 sec after beginning of mec application. (E) Normalized pulsing rate as a function of time since mec application (grey arrow), averaged from *n* = 12 lLNvs (blue) and *n* = 14 sLNvs (pink). The pulsing rate for each neuron was normalized to its value at the beginning of mec application. Shaded regions are SD of the pulsing rate.

**Figure S6.**
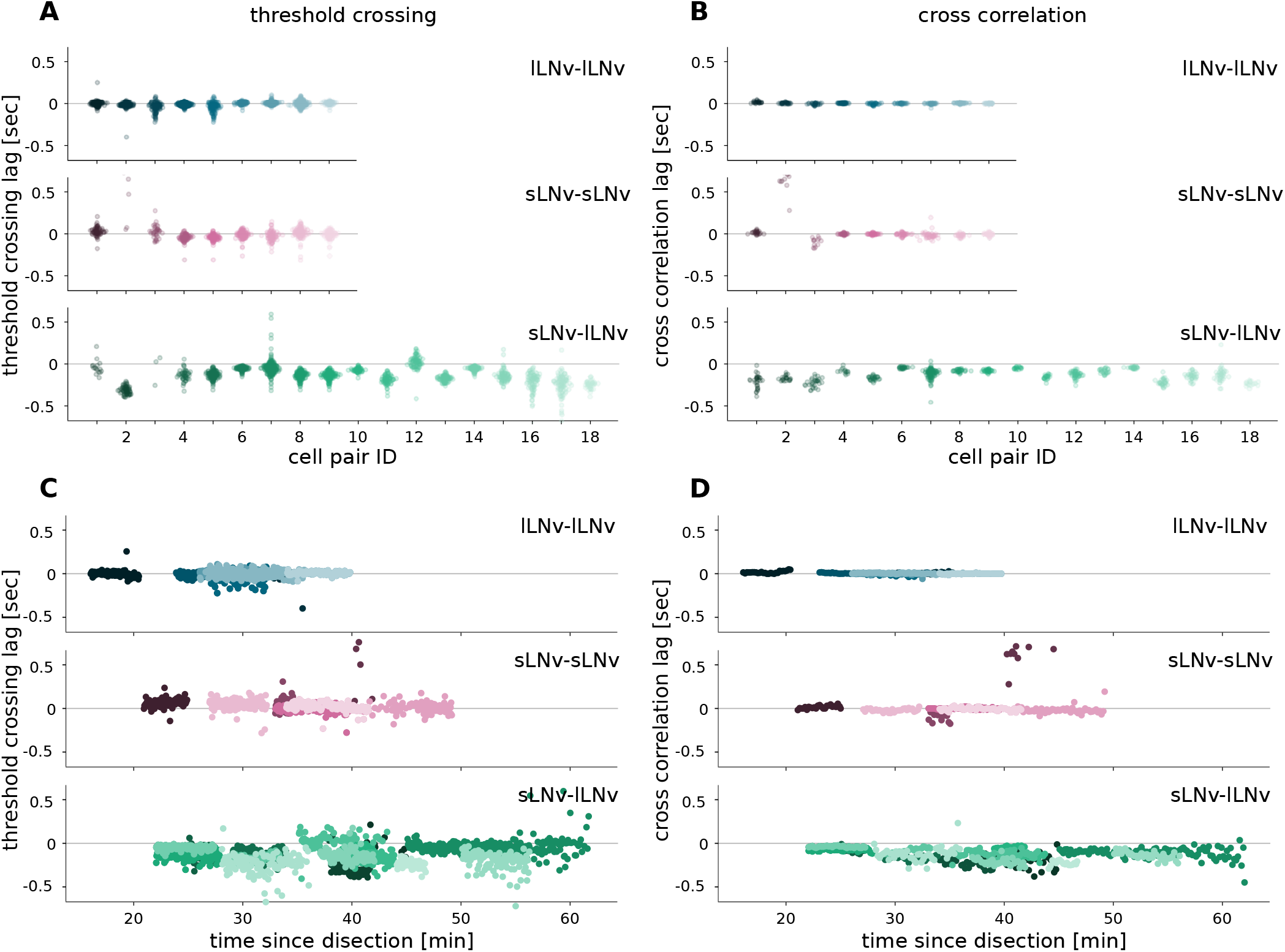
Lags of individual pairs are consistent with population level results. (A-D) Lag of different neuronal pairs obtained from dual recordings by (A, C) threshold crossing analysis and (B, D) local cross-correlation, for lLNv pairs (dots in shades of blue), sLNv pairs (dots in shades of pink) and lLNv-sLNv pairs (dots in shades of green).

**Figure S7.**
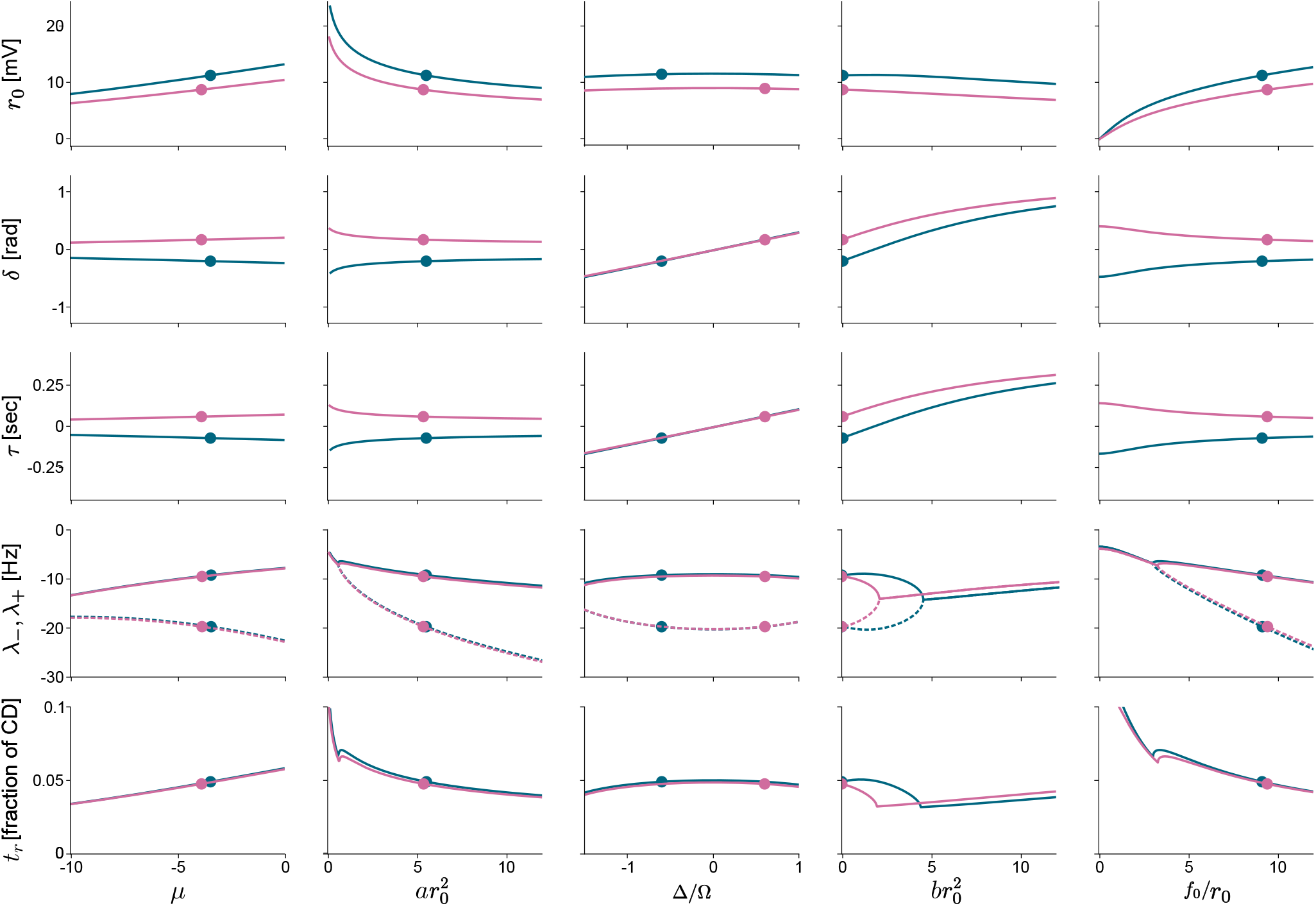
Parameters control different aspects of observables in the theory. Half amplitude *r*_0_, phase lag is *δ*, time lag *τ*, eigenvalues *λ*_+_ (solid) and *λ*_*−*_ (dashed) and relaxation time is *t*_*r*_, are plotted as a function of parameters *µ, a*, Δ, *b* and *f*. Parameters are scaled to have units of sec^*−*1^ to make them comparable, except for detuning Δ which is scaled by the forcing frequency Ω, such that Δ = 1 corresponds to *ω* = 0. Observables are calculated for lLNv (blue dots) and sLNv (pink dots) parameter values (Table I).

